# Venous activation of MEK/ERK drives development of arteriovenous malformation and blood flow anomalies with loss of Rasa1

**DOI:** 10.1101/2021.07.21.453236

**Authors:** Jasper Greysson-Wong, Rachael Rode, Jae-Ryeon Ryu, Kristina D. Rinker, Sarah J. Childs

## Abstract

Vascular malformations develop when growth pathway signaling goes awry in the endothelial cells lining blood vessels. Arteriovenous malformations (AVMs) arise where arteries and veins abnormally connect in patients with loss of RASA1, a Ras GTPase activating protein, and, as we show here, in zebrafish rasa1 mutants. Mutant fish develop massively enlarged vessels at the connection between artery and vein in the tail vascular plexus. These AVMs progressively enlarge and become filled with slow-flowing blood and have a greater drop in pulsatility from the artery to the vein. Expression of the flow responsive transcription factor klf2a is diminished in rasa1 mutants, suggesting changes in flow velocity and pattern contribute to the progression of vessel malformations. Migration of endothelial cells is not affected in rasa1 mutants, nor is cell death or proliferation. Early developmental artery-vein patterning is also normal in rasa1 mutants, but we find that MEK/ERK signaling is ectopically activated in the vein as compared to high arterial activation seen in wildtype animals. MEK/ERK signaling inhibition prevents AVM development of rasa1 mutants, demonstrating venous MEK/ERK drives the initiation of rasa1 AVMs. Thus, rasa1 mutants show overactivation of MEK/ERK signaling causes AVM formation, altered blood flow and downstream flow responsive signaling.

**Summary:** The zebrafish model of *RASA1* capillary malformation and arteriovenous malformation (CM-AVM1) develops cavernous vascular malformations driven by ectopic MEK/ERK signaling in the vein, disrupting flow and downstream mechanosensitive signaling.

## Introduction

The vascular tree relies on an orderly branched structure of progressively sized vessels to effectively transport nutrients and oxygen to cells. Vascular malformations such as arteriovenous malformations (AVMs), angiomas, hemangiomas, aneurysms, and vascular tumors are a result of altered developmental vascular signaling that disrupt the tree-like structure of the vascular system. Capillary malformation-arteriovenous malformation (OMIM: 608354; CM-AVM1) is caused by mutations in the RASA1 GTPase Activating Protein (Eerola et al., 2003). RASA1 is clearly important for vascular development across species as loss of *Rasa1* in mice and *rasa1* knockdown in zebrafish leads to disordered vasculature (Henkemeyer et al., 1995; Kawasaki et al., 2014; Lubeck et al., 2014).

The most prominent presentation of human CM-AVM are capillary malformations (CMs), which appear in ∼95% of patients (Duran et al., 2018; Heuchan et al., 2013; Lapinski et al., 2018; Revencu et al., 2008). CMs are cutaneous beds of permanently dilated capillaries that appear as a purple-red or port-wine ‘stain’ on the skin. About a third of patients have an AVM that directly shunts blood between arterial and venous systems, by-passing capillary beds that normally intercede the two systems. AVMs are fragile, prone to rupture and difficult to treat. The localized nature of these vascular malformations appears to be the result of a somatic second hit that is permissive of lesion formation (Lapinski et al., 2018). Although it is a ubiquitously expressed gene, RASA1 function is necessary in endothelial cells for vascular homeostasis, and the GAP domain is critical for its function (Henkemeyer et al., 1995; Lapinski et al., 2012; Lubeck et al., 2014). Taken together, RASA1’s GAP function within the endothelium is critical for vascular tree development and loss of its activity drives the development of vascular defects leading to CM-AVM.

The signaling pathway upstream of RASA1 has become clearer through genetic analysis. In zebrafish, *rasa1a* morpholino knockdown has similar vascular defects to knockdown of the EphB4 kinase (*ephb4a* morphants) including vessel enlargement in the caudal venous plexus (CVP), lack of caudal blood flow and overabundance of intersegmental veins (ISVs) at the expense of intersegmental arteries (ISAs) (Kawasaki et al., 2014). Similarly, loss of EPHB4 in mice leads to vascular malformations and, in humans, a strikingly similar disease CM-AVM2 caused by mutations in human EPHB4 (OMIM: 618196) suggesting that interactions between the EPHB4 kinase and RASA1 downstream lead to similar endothelial disruption (Amyere et al., 2017; Gerety and Anderson, 2002; Gerety et al., 1999). RASA1 binds EPHB4 in vitro in cultured cells (Kawasaki et al., 2014) and given the importance of EPHB4 in determining vein identity in mice (Amyere et al., 2017; Gerety and Anderson, 2002; Gerety et al., 1999), it is reasonable to hypothesize that arteriovenous identity could be affected with loss of of RASA1 and lead to arteriovenous malformation.

Downstream of RASA1, Ras signaling can activate two downstream pathways, either MEK/ERK or PI3K/AKT/mTORC, both of which are key in arteriovenous specification; each has evidence of being overactivated in different forms of AVM (Alsina-Sanchis et al., 2018; Chen et al., 2012; Chen et al., 2019; Fischer et al., 2004; Fish and Wythe, 2015; Fish et al., 2020; Hong et al., 2006; Iriarte et al., 2019; Kawasaki et al., 2014; Lawson et al., 2001; Lawson et al., 2002; Lubeck et al., 2014; Nikolaev et al., 2018; Ola et al., 2016; Shutter et al., 2000; Wythe et al., 2013; You et al., 2005). RASA1 mutant mice hemorrhage and edema are reversed by inhibition of MEK1/2 (Chen et al., 2019), while zebrafish *rasa1a* morphant fish show fewer venous intersegmental vessels after inhibition of PI3K/mTORC (Kawasaki et al., 2014). However, it remains unclear which pathways downstream of RASA1 drive AVM formation since previous RASA1 loss of function models have not characterized AVMs.

Here, we use a zebrafish model of Rasa1 CM-AVM to understand the real-time development of vascular malformations, and their effect on blood flow and signaling. *rasa1* genetic mutants develop AVMs in the CVP as early as 30 hours post fertilization (hpf). The AVMs progressively enlarge, disrupting blood flow and filling with stagnant blood. Loss of Rasa1 is lethal by 10dpf. We observe that both blood flow velocity and pulsatility are affected by the cavernous malformation, resulting in slower flow in the AVM and a substantial drop in pulsatility from the dorsal aorta to the caudal vein. Correspondingly, expression of the flow responsive transcription factor *klf2a* is diminished, suggesting the presence of the AVM results in changes in flow velocity and pattern, potentially contributing to the progression of vessel malformations through changes in mechanosensory signaling. We see preferential activation of pERK in the vein of *rasa1* mutants and rescue of blood vessel patterning with the inhibition of MEK/ERK signaling. This shows that aberrant venous MEK/ERK signaling is critical in the initiation of Rasa1 AVMs.

## Results

### *rasa1* mutation leads to cavernous AVM development in the tail plexus

There are two RASA1 orthologs in zebrafish. To create a CM-AVM1 model, we used CRISPR-Cas9 to create *rasa1a*^*ca35*^ and *rasa1b*^*ca59*^ mutants (Figure 1-figure supplement 1). Single *rasa1a or rasa1b* mutants are homozygous viable and have mild phenotypes (Figure 1-figure supplement 2) including small ectopic shunts in *rasa1a* mutants, directly between the dorsal aorta (DA) and caudal venous plexus (CVP) at 30hpf that resolve by 48hpf. As a result of the subtle, resolvable vascular phenotype of single mutants, *rasa1a*-/-;*rasa1b-/-* double mutants (hereafter *rasa1*-/- or *rasa1* mutant, generated from *rasa1a-/-;rasa1b+/-* incrosses) were used to characterize all vascular phenotypes.

The CVP of the zebrafish tail is homologous to mammalian vascular plexi where a surplus of vessels develops initially and is gradually refined into an efficient vascular network. Patterning of vessels in the plexus does not follow a strict pattern in contrast to other highly studied beds like the intersegmental vessels in zebrafish. We used confocal microscopy to characterize vessel structure at two key stages of development. 30hpf is a critical point in CVP development since the caudal vein (CV) undergoes angiogenic sprouting between 24hpf and 30hpf. By 30hpf, the CVP has expanded ventrally and blood circulation within the vessel bed is robust. Between 30hpf and 48hpf, pruning and remodeling of the CVP occurs; thus, defects in vessel remodeling would likely become evident by 48hpf. To detect the AVMs, we measured the diameter of the largest vein in the CVP. *rasa1* mutants show vein enlargement (≥1.5x average largest vein in wildtype) as early as 30hpf (*rasa1-/-:* 86%±25 penetrance vs. WT: 8%±14, p=0.0050, Figure 1A, E-G, N, P), averaging 73.5µm±29.0 in diameter whereas wildtype largest veins were 43.5µm±21.6 (LV: p=0.0018, Figure 1A-D, N, P). At 48hpf, the developing cavernous AVM is consistently located at the posterior of the tail plexus, connecting the rostral DA to the rostral CVP, subsuming a portion of the capillaries in the CVP. The largest vein in mutants at 48hpf measures 108.4µm±70.8 whereas wildtype veins narrow to 22.3µm±4.4 (LV: p<0.0001, Figure 1H-M, P; *rasa1-/-*: 69%±3 penetrance vs. WT: 0%, p=0.02; Figure 1N). *rasa1* mutant embryos develop severe edema by 5dpf and lethality by 10dpf (Chi-sq: p<0.0001, Figure1-Figure Supplement 1). As a control, we measured the average DA diameter upstream of the malformation. The DA does not differ in size between wildtype and mutants at either timepoint (Figure 1A-M, O). At 30hpf, wildtype DA diameter is 17.5µm±3.2 versus *rasa1-/-* at 17.4µm±3.2 (p>0.99) and, by 48hpf, the DA widens to 21.4µm±3.2 in wildtypes and 21.4µm±4.5 in mutants (p>0.99). To demonstrate the significant vessel enlargement in mutants, we used Simpleware for 3D rendering of the vessel morphologies in wildtypes and *rasa1* mutants (Figure 1D, G, J, M-N).

**Figure 1.**
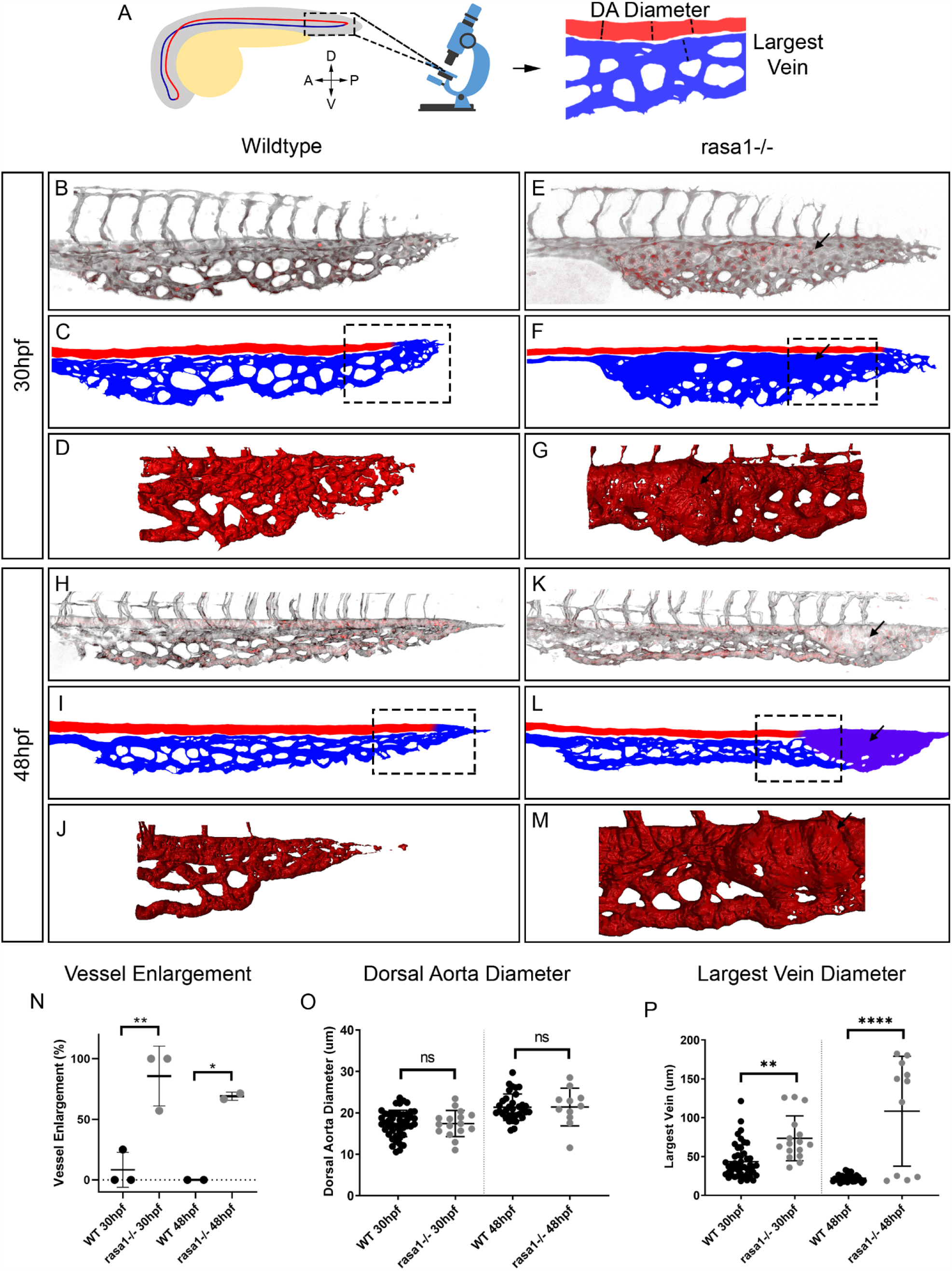
Confocal imaging of *rasa1-/-* illustrates that vessel enlargement developing in the caudal venous plexus by 30hpf without affecting the dorsal aorta. **A:** Diagram showing the location of the caudal venous plexus, with the boxed area illustrating where confocal images were taken as well as the orientation. A=anterior, P=posterior, D=dorsal, V=ventral. **B:** Diagram showing how dorsal aorta (DA) diameter was measured in triplicate and how largest vein was measured. **B, E, H, K:** Confocal images of the caudal venous plexus of wildtype and *rasa1* mutants on *Tg(flk:EGFP;gata1a:dsRed)*. Black is the *flk:EGFP* endothelium, red is *gata1a:dsRed* red blood cells. **C, F, I, L:** Schematic of the vessel structures showing red is artery, in this case the dorsal aorta and blue is the venous structure, the caudal venous plexus and purple, marking vessels of unknown arteriovenous identity. **D, G, J, M:** Simpleware was used on high resolution confocal images to create 3D renderings of the flow return, where malformations in *rasa1* mutants develop. Note boxes in C, F, I, L only illustrate approximate location of the 3D renderings, with the 3D renderings being generated from high resolution images of different embryos. **N-O:** Quantification of confocal images of wildtype (WT) and *rasa1* mutant embryos at 30hpf and 48hpf. **N:** Penetrance of vessel enlargement (≥1.5x average largest wildtype vein diameter) at 30hpf and 48hpf (30hpf: WT n=13, *rasa1-/-* n=16, N=3, p=0.005. 48hpf: WT n=10, *rasa1-/-* n=10, N=2, p=0.024.) **O:** The averaged wildtype DA diameter was not significantly different than in mutants at either timepoints (30hpf: WT n=13, *rasa1-/-* n=16, N=3, p>0.99. 48hpf: WT n=12, *rasa1-/-* n=11, N=3, p>0.99). **P:** The largest vein at 30hpf is larger in mutants and is further enlarged at 48hpf (30hpf: WT: n=13, *rasa1-/-:* n=16, N=3, p=0.0018, 48hpf: WT: n=12, *rasa1-/-:* n=11, N=3, p<0.0001). P-values were calculated using a one-way ANOVA with Sidak’s correction for multiple comparison. Error bars represent ±SD.

**Figure 1- figure supplement 1.**
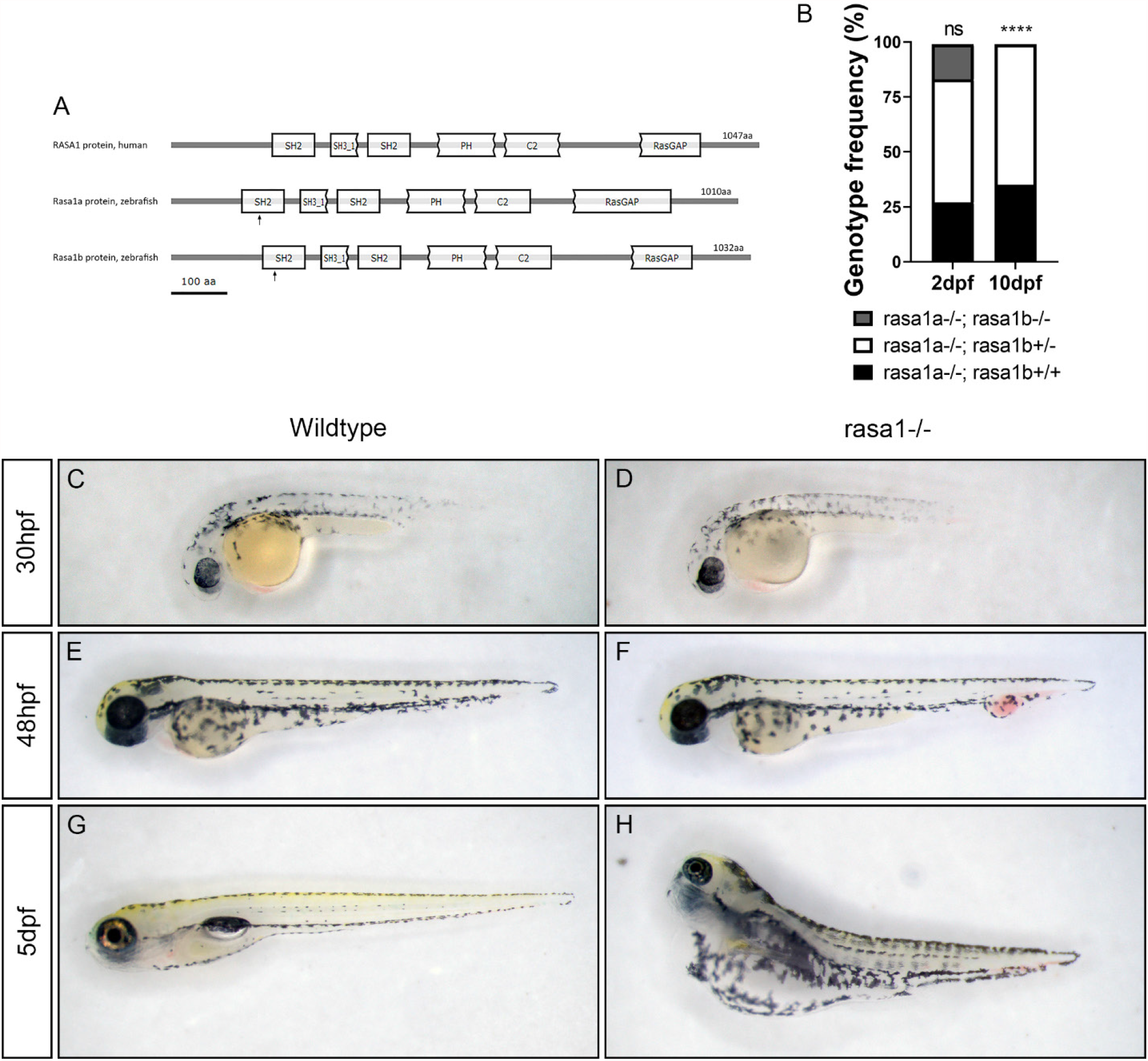
*rasa1* mutants have vascular malformations in the tail vessels by 30hpf and develop severe edema by 5dpf, with *rasa1-/-* being lethal by 10dpf. **A:** Diagram of key protein domains in human RASA1 versus zebrafish Rasa1a and Rasa1b modified from CDvist. Arrows indicate where our *rasa1a*^*ca35*^ and *rasa1b*^*ca59*^ mutant lines are truncated. **B:** Genotyping of clutches at 2dpf and 10dpf reveal that complete loss of *rasa1* is lethal by 10dpf (Chi-sq: p<0.0001). **C-H:** Stereoscope images illustrate the relative size and severity of the vascular malformation in *rasa1-/-* at 30hpf, 2dpf and 5dpf. **C, D:** At 30hpf, *rasa1* mutants are almost indistinguishable to wildtypes under a stereoscope. **E-F:** The vascular malformation observed in the tails of mutants are obvious even under a stereoscope at 2dpf versus wildtype controls. **G-H:** By 5dpf, *rasa1-/-* develop severe edema.

**Figure 1- figure supplement 2.**
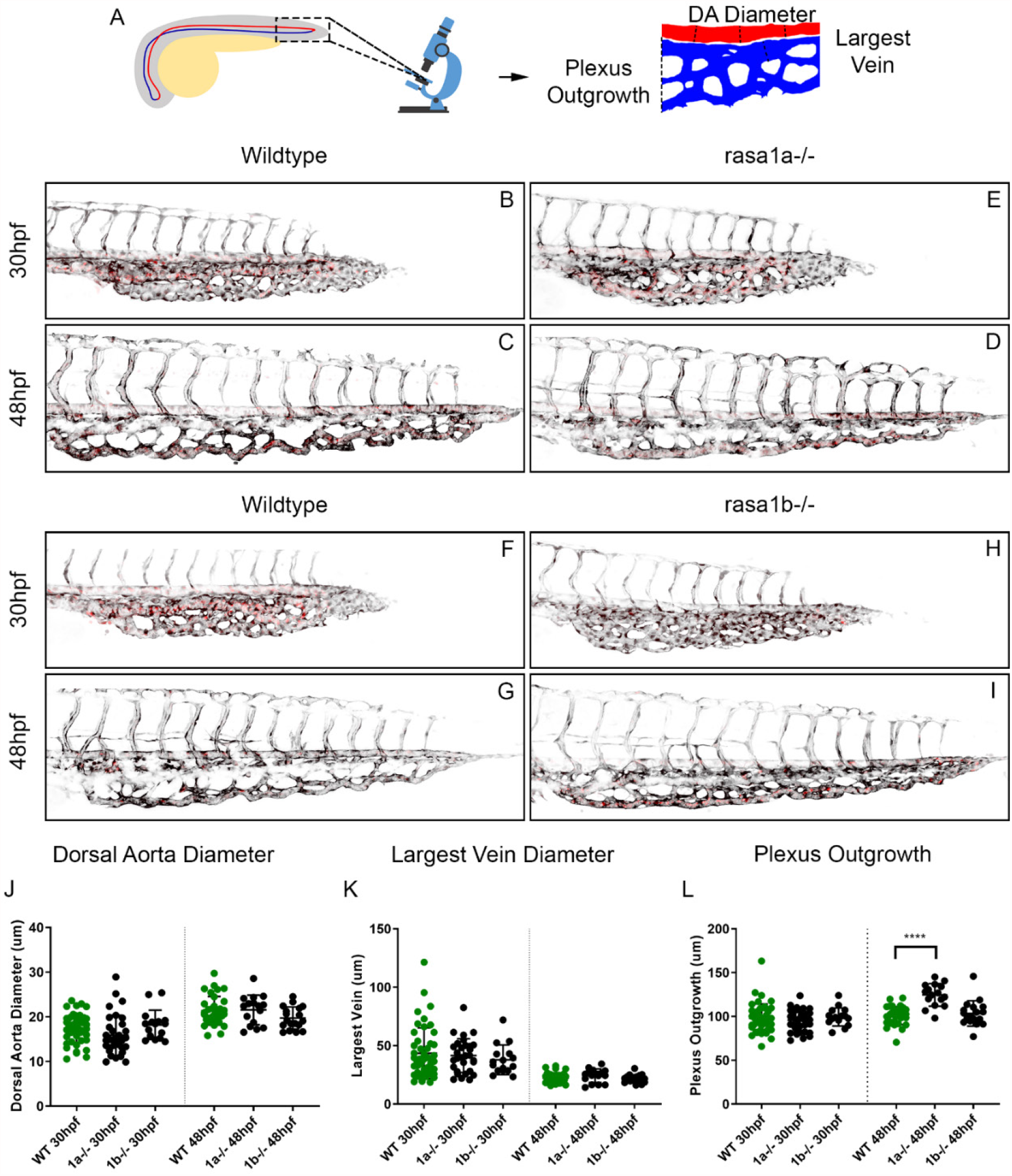
*rasa1a* and *rasa1b* singles mutants have very mild vascular phenotypes. **A:** Schematic illustrates how confocal images were taken of wildtype and *rasa1a-/-* and *rasa1b-/-* embryos at 30hpf and 48hpf. **B-I:** Confocal microscopy of *rasa1a-/-* and *rasa1b-/-* embryos on the *Tg(flk:EGFP;gata1a:dsRed)* background at 30hpf reveal small, infrequent, ectopic connections between the dorsal aorta (DA) and caudal venous plexus which resolve by 48hpf. Black is the *flk:EGFP* endothelium, red is *gata1a:dsRed* red blood cells. 30hpf: *rasa1a*-/-: 30.3%, n=33, wildtype: 17.4%, n=23, N=6; *rasa1b-/-*: 33.3%, n=9, wildtype: 12.5%, n=8, N=2; 48hpf: *rasa1a*-/-: 0%, n=17, wildtype: 0%, n=13, N=2; *rasa1b-/-*: 0%, n=19, wildtype: 0%, n=17, N=3). **J, K, L:** Vessel measurements were quantified, no significant changes were observed in the DA diameter, largest vein diameter or plexus outgrowth at either timepoint between wildtypes and mutants except for *rasa1a-/-* had a significantly more advanced plexus at 48hpf than wildtypes (p<0.0001). P-values were calculated using a one-way ANOVA with Sidak’s correction for multiple comparison. Error bars represent ±SD.

### Vascular malformation alters blood flow up- and down-stream of the lesion

We next determined how the abnormal vascular architecture leads to altered blood flow patterns as this has not been examined in vivo in a *RASA1* model. We determined that heart rate is not changed at either 30hpf or 48hpf, suggesting the heart output is normal (30hpf: p=0.17, 48hpf: p=0.13, Figure 2-figure supplement 1). Using high speed video imaging, we imaged blood flow through the DA and CVP at 30hpf and 48hpf (videos 1-4). Mean velocities were calculated in the DA at the CV proximal to the flow return (Figure 2A, Figure 2-figure supplement 1). Heatmaps of representative single embryos (Figure 2B-E) and averaged from multiple embryos (Figure 2F-I) illustrate consistent flow changes in *rasa1* mutant malformations. We focused on three areas of the vasculature: 1) the DA, which is not expected to differ between mutants and wildtypes as it is upstream of the malformation, 2) the caudal vein, 3) and the point where the DA ‘turns’ 180 degrees into the CVP, which we named the flow return, also the site where the malformation develops in mutants.

**Figure 2.**
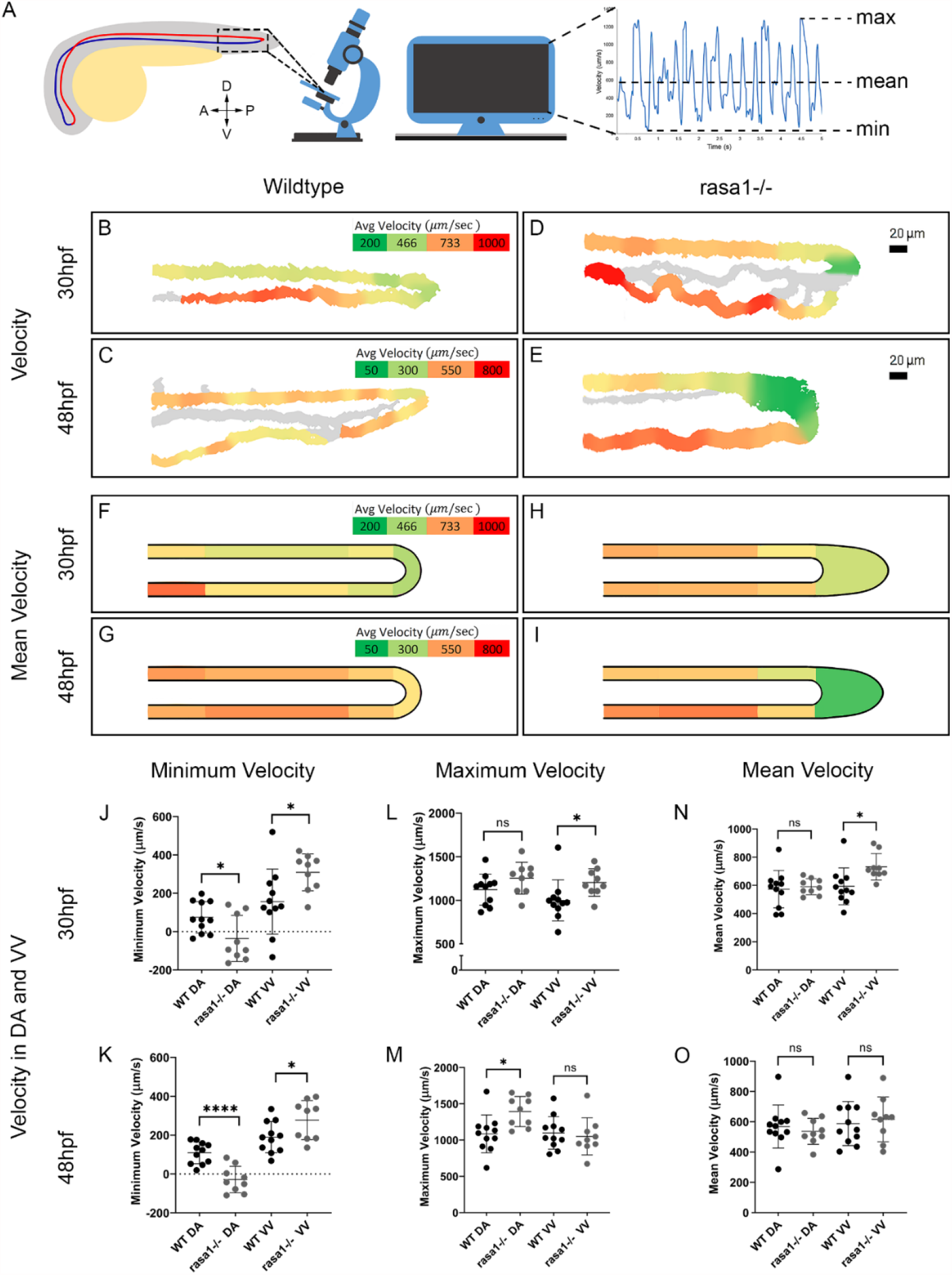
Blood flow velocity is impacted by the vascular malformations in *rasa1-/-* embryos. **A:** Diagram illustrating where videos were taken and how velocity metrics were calculated. **B-E:** Velocity heatmaps of representative wildtype and *rasa1* mutant embryos. **F-I:** Mean velocity heatmaps wildtype (WT) and *rasa1* mutant embryos. **J-O:** Quantification of minimum, maximum and mean velocities in wildtypes and *rasa1* mutants at 30hpf and 48hpf (WT: 30hpf and 48hpf: n=11. *rasa1-/-:* 30hpf and 48hpf: n=9). **J**,**K:** Minimum velocity at 30hpf and 48hpf of wildtype and *rasa1* mutants in the DA and ventral vein (VV). **L**,**M:** Maximum velocity at 30hpf and 48hpf of WT and *rasa1* mutants in the DA and VV. **N**,**O:** Mean velocity at 30hpf and 48hpf of WT and *rasa1* mutants in the DA and VV. P-values were calculated using a one-way ANOVA with Sidak’s correction for multiple comparison. Error bars represent ±SD.

Blood flow rates vary according to the size and location of the vessel and developmental stage. In 30 hpf zebrafish there is typically high velocity flow in the DA (573.1µm/s±132.7), slower flow in the return (483.9µm/s±108.7), and fast flow in the CV (593.2µm/s±130.6, Figure 2B, F, N,Figure 2-figure supplement 1). Thus, in wildtypes the flow velocity in the DA is similar to the CV (p=0.62) at this early stage. In *rasa1* mutants at 30hpf, the average velocity in the DA is 591.1µm/s±56.9, but the CV has a significantly faster velocity than the DA at 731.7µm±93.8 (p=0.0062, paired t-test, Figure 2D, H, N, Supp. 4C). Mutants have a lower average flow speed at the flow return relative to the DA and CV (522.3µm/s±152.6) but did not significantly differ from wildtype (p=0.90, multiple t-tests with Holm-Sidak correction). Thus, while average flow velocity in the DA and return is similar between wildtypes and mutants, there is faster flow in the CV of mutants (DA: p>0.99, CV: p=0.034, one-way ANOVA).

**Figure 2- figure supplement 1.**
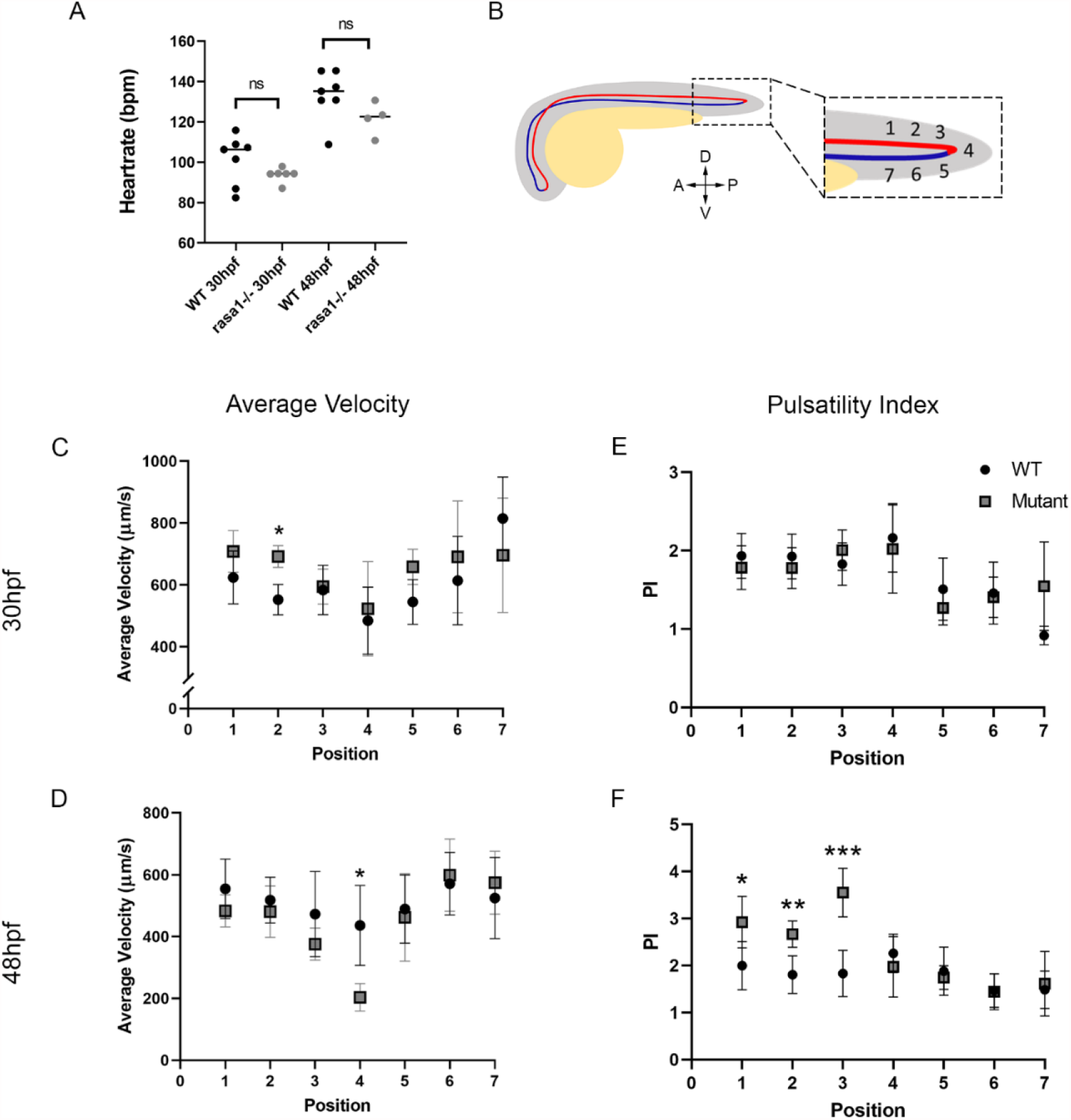
Heart rate is not affected in *rasa1* mutants; average velocity and pulsatility index with positional resolution shows effects of vascular malformations throughout the DA and CVP. **A:** Heart rate in not impaired in *rasa1* mutants versus wildtypes at either 30hpf or 48hpf (HR: 30hpf: WT: 101.3bpm±12.2, n=7, *rasa1-/-*: 93.7±3.6, n=6, p=0.17, N=1. 48hpf: WT: 133.4±12.4, n=7, *rasa1-/-*: 121.7±8.2, n=4, p=0.13, N=1, unpaired t-tests). **B:** Diagram showing the relative location of positional data, with position 1-3 on the dorsal aorta, position 4 at the flow return and position 5-7 running along the ventral vein. **C-D**. Positional data for average velocities. **C:** Average velocities at 30hpf reveal significant differences between WT and *rasa1-/-* at position 2 (WT: 551.8±49.5µm/s, *rasa1*-/-: 691.5µm/s±35.3, p=0.00062). **D:** At 48hpf, average velocities differ at position 4 (WT: 436.3µm/s±129.4, *rasa1*-/-: 203.3µm/s±44.8, p4=0.038). **E-F:** Positional data for PI. **E:** No significant differences are seen in PI at 30hpf. **F:** At 48hpf, pulsatility is significantly different between WT and *rasa1-/-* at position 1 (WT: 2.0±0.51, *rasa1-/-*: 2.9±0.54, p1=0.040), position 2 (WT: 1.8±0.40, *rasa1-/-*: 2.7±0.28, p2=0.0031), position 3 (WT: 1.8±0.49, *rasa1*-/-: 3.6±0.52, p3=0.00014). P-values were calculated for positional data by multiple t-tests with Holm-Sidak correction for multiple comparisons. Error bars represent ±SD.

At 48hpf, wildtype DA and CV have similar average velocities (DA: 568.2µm/s±142.2, CV: 588.2µm/s±145.5, p=0.73) and lower velocity in the flow return (436.3µm/s±129.4, Figure 2C, G, O, Fig 2-figure supplement 1). In *rasa1* mutants at 48hpf, the malformation has enlarged and is now an AVM. However, there are no significant difference between DA or CV velocities (p=0.085). The average flow in the DA is 536.8µm/s±85.8 and in the CV is 616.4µm/s±147.8 with a drop in flow speed at the return (203.3µm/s±44.8, Figure 2E, I, O, Figure 2-figure supplement 1). There is, however, a significant drop in velocity at the flow return in mutants versus wildtypes (WT: 436.3µm/s±129.4, vs. rasa1-/-: 203.3µm/s±44.8, p4=0.038) likely because of a larger malformation.There are no significant changes in DA or CV velocities between wildtypes and mutants (DA: p=0.98, CV: p=0.98).

Due to the pulsatile nature of blood flow, average velocities do not capture the complexity of flow changes that occurs through the cardiac cycle. At 30hpf and 48hpf, during diastole the minimum velocities in wildtypes are not significantly different the DA (30hpf: 74.4µm/s±82.0, 48hpf: 109.1µm/s±55.9) or CV (30hpf: 156.3µm/s±169.8, p=0.42, 48hpf: 189.7µm/s± 80.6, p=0.075, Figure 2J-K). In contrast, at 30hpf in *rasa1* mutants, there is a drastic difference in minimum velocities between the two vessels with the DA minimum velocity at - 35.9µm/s±120.5 versus the CV at 309.0µm/s±96.3 (p<0.0001, Figure 2J). A negative number indicates backflow in the DA. At 48hpf, the same trend is observed, with the minimum velocity in the DA (−28.6µm/s±67.9) being significantly lower than the CV (277.2µm/s±100.7, p<0.0001, Figure 2K). In comparing *rasa1* mutants to wildtypes, the minimum DA velocity is significantly lower in mutants than wildtypes at both 30hpf and 48hpf (30hpf: p=0.026; 48hpf: p<0.0001) but is higher in the mutant ventral vein at both timepoints (30hpf: p=0.028, 48hpf: p=0.045, Figure 2J-K).

We also examined maximum velocities during systole. In wildtypes, we find no significant difference in velocities at 30hpf or 48hpf (30hpf: p=0.46, 48hpf: p>0.99, Figure 2L-M). At 30hpf, *rasa1* mutants maximum velocities were not different between the two vessels (p=0.97, Figure 2L). When we compare *rasa1* mutants to wildtypes at 30hpf, the maximum velocity is not significantly different in the DA or CV (DA: p=0.45, CV: p=0.089). By 48hpf, *rasa1* mutants have an increase in DA maximum velocity (1394µm/s±207.9) over the CV (1052µm/s±256.5, p=0.017, Figure 2M). Comparing wildtypes to mutants shows the maximum velocity in the DA of *rasa1* mutants (1394µm/s±207.9) is significantly higher than wildtypes (1086µm/s±260.9, p=0.027) with no significant difference in the CV (p=0.99).

Taken together, the vessel malformation and AVM affect velocity extremes. Over both timepoints, an increase in minimum velocity in the CV of mutants is paired with a decrease in the minimum velocity in DA. This suggests the malformation creates flow velocity patterns that are very different to what normal vessels would experience where an increase in flow in one vessel type corresponds to a decrease in flow in the other vessel type.

### Increased drop in pulsatility between the dorsal aorta and caudal vein in *rasa1* mutants

Pulsatility from discrete heart beats is reflected in the amplitude of change from maximal blood velocity to the minimum velocity (Figure 3A-B, Eq. 1). We created representative single embryo pulsatility heatmaps (Figure 3C-F), and averaged pulsatility heatmaps from 7 embryos (Figure 3G-J) to demonstrate variation in flow pulsatility across the DA and CVP. In wildtypes at 30hpf, heatmaps reveal higher pulsatility in the DA (1.9±0.3) and lower pulsatility in the CVP (1.4±0.2, p=0.0006, Figure 3C, G, K) as we would expect for a vessel at a distance from the heart. By 48hpf, we see that flow pulsatility evens out across the two vessels, with the DA (1.7±0.3) not statistically different than in the ventral vein (1.6±0.2, p=0.68, Figure 3E, I, M). In contrast, 30hpf *rasa1* mutants have elevated pulsatility in the DA (2.2±0.3) relative to the ventral vein (1.2±0.1, p<0.0001, Figure 3D, H, K). By 48hpf, pulsatility in the DA remains elevated relative to the CV (DA 2.7±0.5 and CV 1.3±0.3, p<0.0001, Figure 3F, J, M). At both 30hpf and 48hpf, flow in the DA of mutants is more pulsatile that in wildtypes (30hpf: p=0.023, 48hpf: p<0.0001) with no change in CV pulsatility (30hpf: p=0.23, 48hpf: p=0.19, Figure 3C-J, K, M).

**Figure 3.**
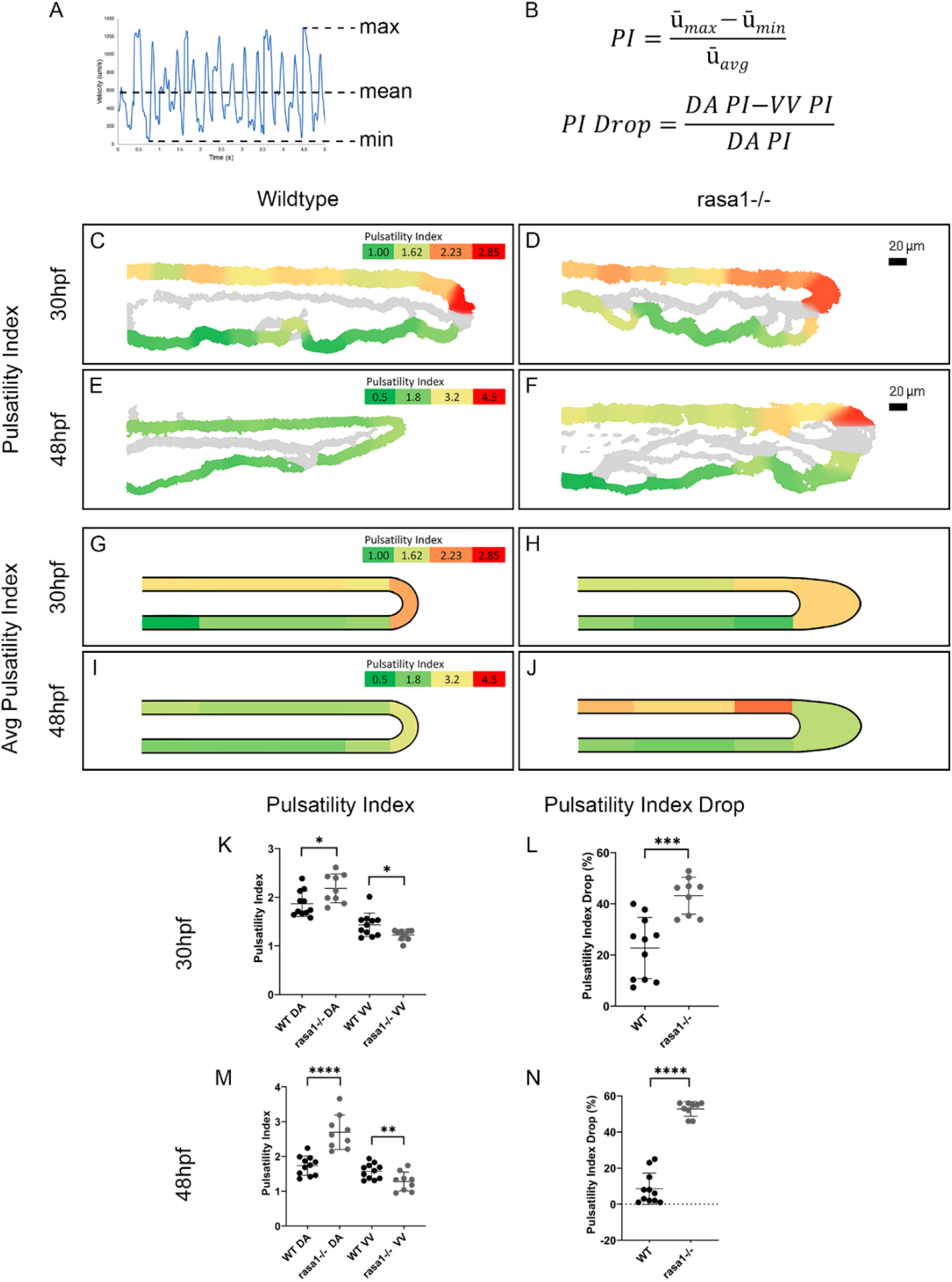
Blood flow pulsatility is impacted by vascular lesions in *rasa1* mutants. **A, B:** Diagram and equations illustrating how pulsatility index (PI) and PI drop were calculated. **C-F:** PI heatmaps of representative wildtype and *rasa1* mutant embryos. **G-J:** Average PI heatmaps wildtype and *rasa1* mutant embryos. **K-L:** PI at 30hpf and 48hpf, respectively, of WT and *rasa1* mutants in the DA and VV. **M, N:** PI drop at 30hpf and 48hpf, respectively, of WT and *rasa1* mutants in the DA and VV. P-values were calculated using a one-way ANOVA with Sidak’s correction for multiple comparison. Error bars represent ±SD.

We next mapped flow pulsatility positionally over three locations in the DA (positions 1-3), in the flow return (position 4) and three locations in the CV (positions 5-7) (Figure 1-figure supplement 1). At 30hpf, there is a stereotypical pattern of flow pulsatility in wildtypes with more pulsatile flow at the 3 DA positions as well as the flow return, and then an immediate drop in pulsatility across its entire length of the CV. At 30 hpf there are no positional differences in wildtypes and mutants. At 48hpf, pulsatility is consistent across the DA, flow return and CV in wildtypes. However, in mutants there is an increase in PI at 48hpf in the 3 DA positions (positions 1-3, p1=0.040, p2=0.0031, p3=0.00014, Suppl. Figure 3F).

The drop in pulsatility from the DA to the ventral vein of the CVP (Figure 3A-B, Eq. 2) results from the effect of the malformation on artery and vein flow. At 30hpf, the average drop in PI from the DA to the VV in wildtypes is 22.7±12.0 compared to 43.2±7.2 in *rasa1* mutants (p=0.0003). By 48hpf, the drop in PI is reduced in wildtypes (8.5±8.7), but more drastically elevated in mutants (52.8±4.1, p<0.0001). This indicates that as the malformation expands, pulsatility is more severely impacted.

Since we detected large changes in velocity and pulsatility, it follows that mechanosensation by endothelial cells and regulation of downstream flow responsive genes could be altered in *rasa1* mutants. Using in situ hybridization, we visualized expression of *klf2a*, a transcription factor that is downregulated by flow in the trunk, at 56hpf. Wildtype embryos have strong *klf2a* staining in both the artery and vein of the trunk. *rasa1* mutants have lower *klf2a* staining in the trunk, which ends more anteriorly (as indicated with arrows; Figure 1-figure supplement 1).

Together these data demonstrate blood flow is strongly altered in the *rasa1* AVM, affecting the upstream input and downstream output vessels surrounding the AVM. This leads to a greater pulsatility and changes to mechanosensory signaling reflected in gene expression of *klf2a*.

### AVMs in *rasa1* mutants do not result from changes in endothelial cell number or migration

**Figure 3- figure supplement 1.**
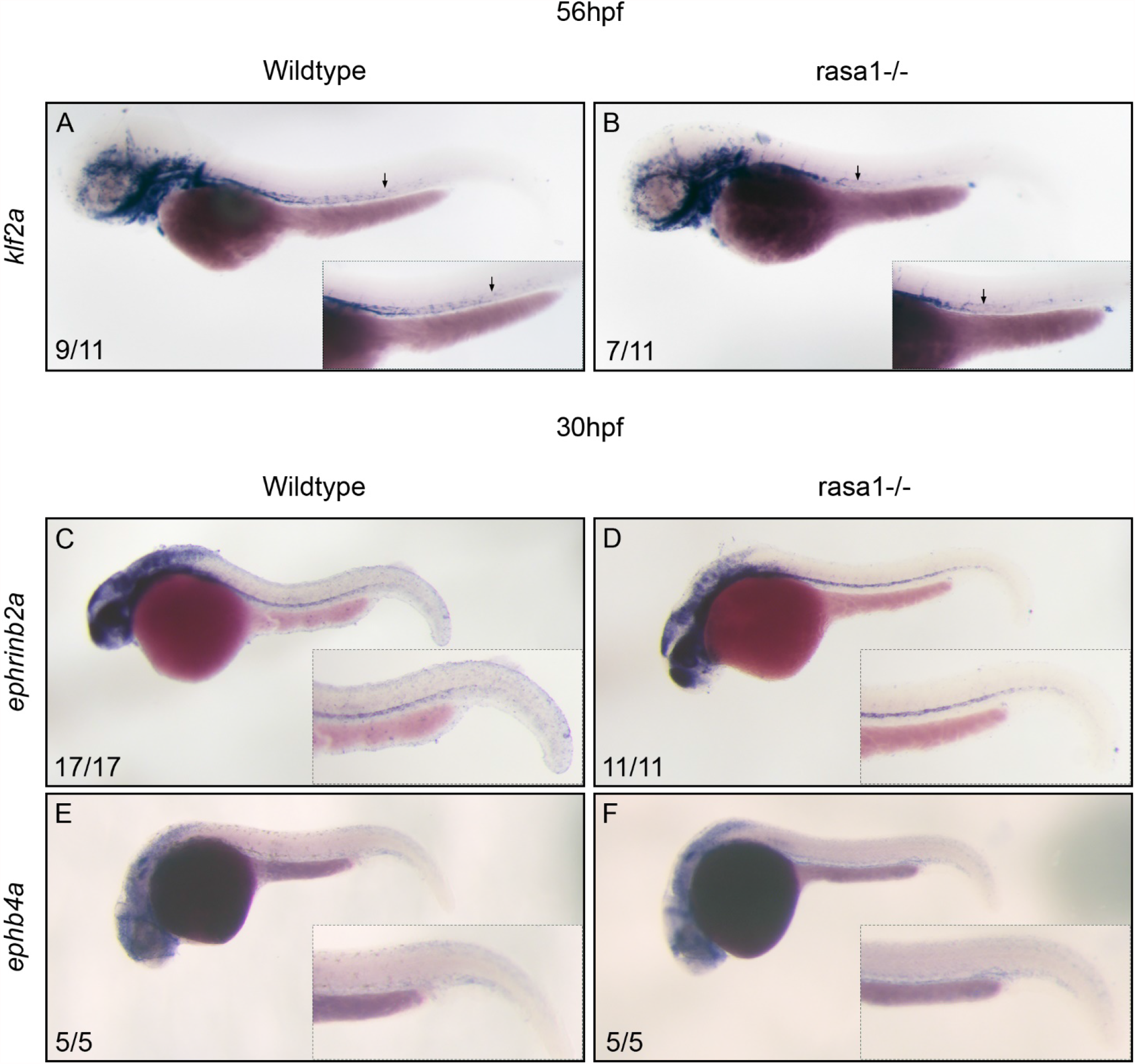
*ephrinb2a* and *ephb4a* ISH reveal no changes in early arteriovenous specification but flow responsive *klf2a* staining shows decreased expression in *rasa1* mutants. In situ hybridization in wildtype and *rasa1* mutants (**A-B):** *klf2a* shows a decrease in staining intensity above the yolk extension at 56hpf in mutants. Arrows indicate the posterior-most position of consistent *klf2a* staining. **(C-D)** Arterial *ephrinb2a* and venous *ephb4a* (**E-F**) in wildtypes and *rasa1-/-* show no obvious difference in staining at 30hpf.

AVMs could develop from an over-proliferation of endothelial cells or a collapse of a plexus into a singular vessel due to apoptosis. However, we find no change in total endothelial cell number between wildtypes and mutants at either timepoint (30hpf: p=0.99, 48hpf: p>0.99, Figure 4K). Proliferation, as detected by phospho-histone H3 (PHH3) staining and show no significant change at 30hpf or 48hpf in mutants versus controls (30hpf: p=0.98, 48hpf: p>0.99, Figure 4A-E). We saw no change in apoptotic cells using immunostaining for cleaved caspase 3 (30hpf: p>0.99, 48hpf: p>0.99, Figure 4F-J). Overall, this suggests that malformations in *rasa1* mutants do not arise from differences of endothelial cell number.

**Figure 4.**
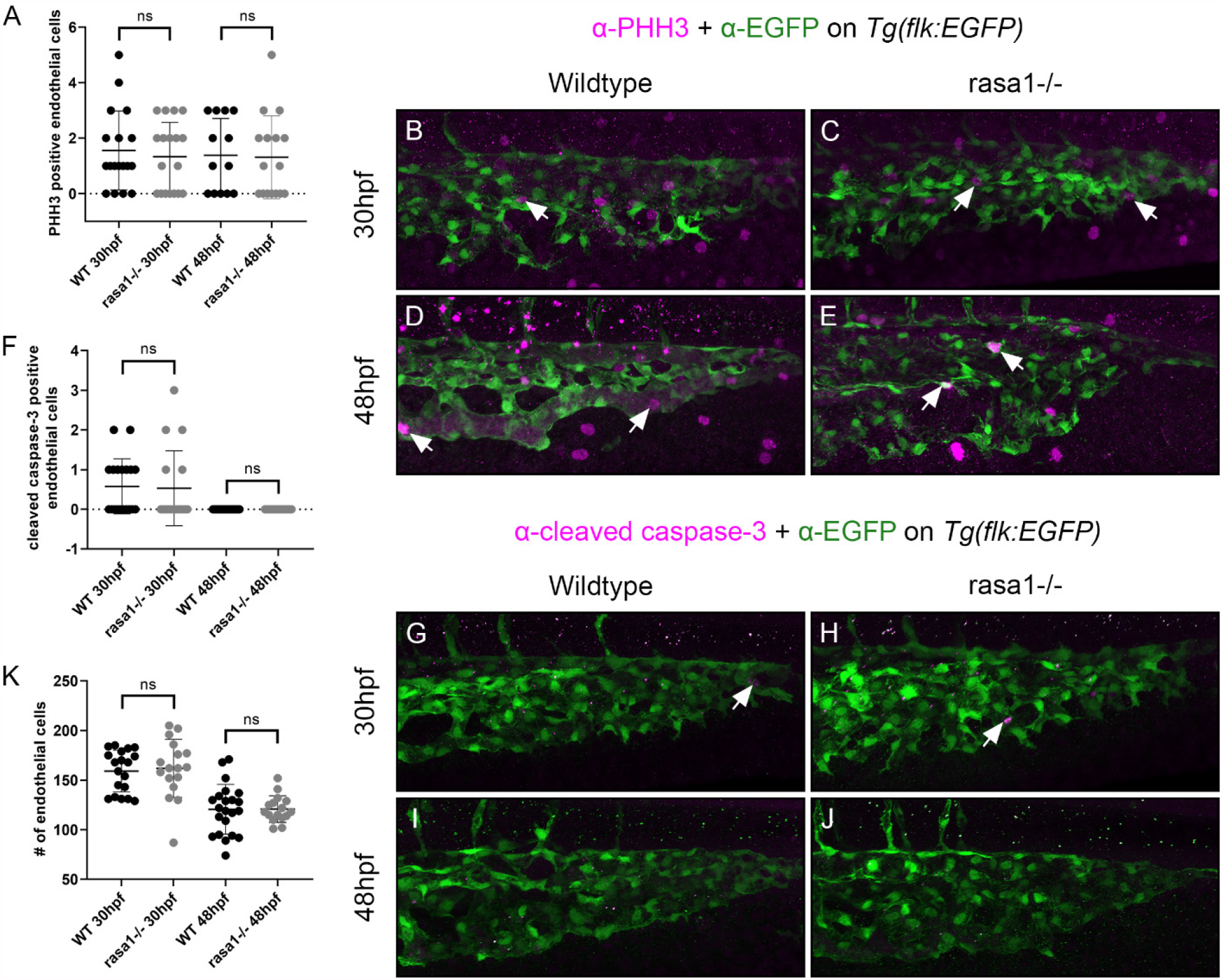
Proliferation and cell death do not drive AVM formation in *rasa1-/-*. **A-E:** Antibody staining of proliferative cells marked by phospho-histone H3 (PHH3) and **F-J:** apoptotic cells with cleaved caspase-3 (cc3) was performed quantified as well as **K:** endothelial cell numbers in the tail vessels based on their transgenic background, *Tg(flk:EGFP)*. **A:** PHH3 staining showed no significant elevation at 30hpf or 48hpf versus controls (30hpf: WT: 1.6cells±1.4, n=18, *rasa1*-/-: 1.3cells±1.2, p=0.98, n=18, N=2. 48hpf: WT: 1.4cells±1.3, n=13, *rasa1*-/-: 1.3cells±1.5, p>0.99, n=16, N=2). **B-E**. Confocal images show PHH3 staining and arrows indicate PHH3+ endothelial cells in wildtypes and mutants at both timepoints. **F**. Apoptotic cells marked by cc3 also did not show any differences between wildtypes and mutants for either timepoint (30hpf: WT: 0.6cells±0.7, n=19, *rasa1*-/-: 0.5cells±0.9, p>0.99, n=17, N=2, 48hpf: WT: 0cells±0, n=22, *rasa1*-/-: 0cells±0, p>0.99, n=17, N=2). **G-J**. Confocal images show cc3 staining and arrows indicate cc3+ endothelial cells in wildtypes and mutants at 30hpf. No cc3 staining in the endothelium was observed at 48hpf. **K**. Endothelial cell counts reveal no significant difference in cell number between wildtypes and mutants at either timepoint (30hpf: WT: 159.2cells±21.1, n=19, *rasa1*-/-: 161.9cells±29.3, *rasa1-/-*: n=17, p=0.99, N=2, 48hpf: 48hpf: WT: 120.6cells±25.1, n=22, *rasa1*-/-: 120.8cells±13.4, *rasa1-/-*: n=17, p>0.99, N=2). P-values were calculated using a one-way ANOVA with Sidak’s correction for multiple comparison. Error bars represent ±SD.

Differences in endothelial behavior might underlie vascular malformation development and progression. For instance, failed sprouting and migration of endothelial cells could result in the formation of vascular malformations. Thus, we quantified the migration distance of wildtype and *rasa1* CVP endothelial cells by imaging the CVP at key developmental windows and CVP migration speed through timelapse imaging. Measuring the furthest extent of the CVP from the DA at different stages, we find no difference in CVP migration distance at 30hpf (WT: 98.0µm±16.3, vs. *rasa1*-/-: 101.7µm±16.7, p=0.89, Figure 4-figure supplement 1). However, the CVP is significantly larger at 48hpf in mutants than wildtypes (WT: 101µm±10, vs. *rasa1*-/-: 128µm±26, p<0.0001, Figure 4-figure supplement 1) that might reflect the swelling of the AVM and not true migration. There is no change in mutant migration speed from timelapse imaging between 24–30hpf, a critical window in CVP development as the CV actively sprouts to form the CVP (WT: 1.7µm/hr±2.0, *rasa1-/-:* 0.99µm/hr±2.1 (p=0.48; Figure 4-figure supplement 1, videos 5-6).

*rasa1* is expressed ubiquitously so we tested whether AVMs could be found in another venous vascular plexus. We imaged the subintestinal venous plexus (SIVP), that develops slightly later, and expands between 58hpf and 76hpf over the surface of yolk sac (Goi and Childs, 2016). We find no obvious vessel malformations or change in SIVP migration distance at 58hpf (p=0.45, Figure 4-figure supplement 1) or at 76hpf (p=0.20). Overall, these results indicate that there is no substantial impairment of endothelial migration in *rasa1* mutants. No AVMs were observed in the animals in other locations including the cerebral circulation. Thus, the CVP appears particularly sensitive to AVM development after loss of *rasa1*.

### Overactivation of venous MEK/ERK signaling in developing vascular malformations

Upregulation of MEK/ERK signaling is observed in vascular anomalies including mouse and human RASA1 mutant cells. pERK immunostaining, a readout of active MEK/ERK signaling, reveals fewer pERK positive nuclei in the DA of *rasa1* mutants at 30hpf (1.6cells±1.4) versus wildtypes (4.3cells±4.3, p=0.03, Figure 5A-B). In contrast, we find a significant increase in pERK nuclei in the vein with 2.0 ±2.6 pERK positive cells in wildtype versus 5.1±2.9 cells in mutants (p=0.0020). There is no change in pERK positive nuclei in the intersegmental arteries (p=0.52) sprouting from the DA (intersegmental veins have yet to sprout from the CVP at 30hpf).

**Figure 4- figure supplement 1.**
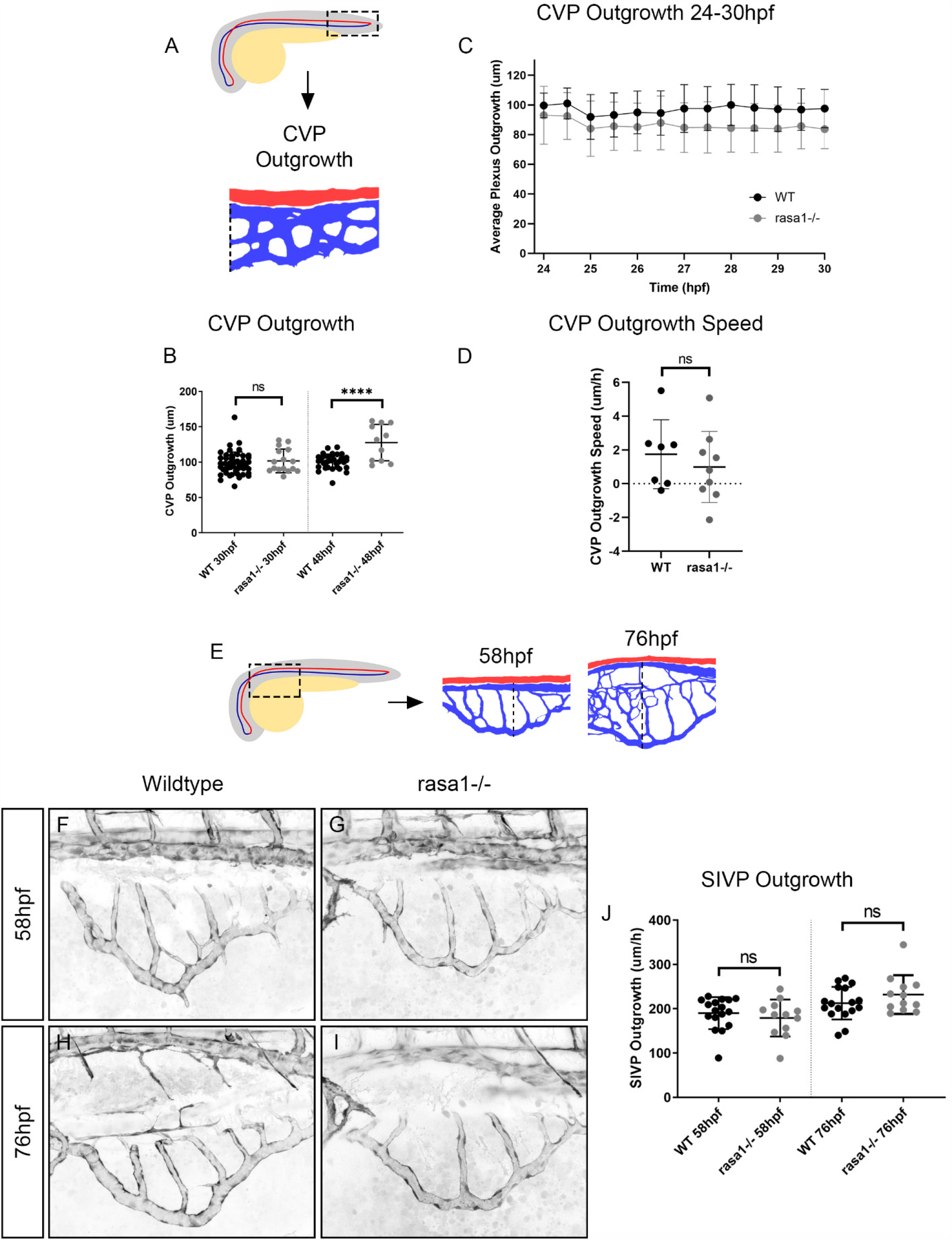
Endothelial migration is unchanged in *rasa1* mutants. **A:** Diagram illustrating how CVP outgrowth was measured, perpendicularly from the DA to the ventral-most aspect of the CVP. **B:** Plexus outgrowth is unchanged between wildtypes and mutants at 30hpf (p=0.89) but is more advanced at 48hpf in mutants than wildtypes (p<0.0001). **C**,**D:** There is no change in *rasa1* mutant migration speed from 24–30hpf from 1.7µm/hr±2.0 in wildtypes to 0.99µm/hr±2.1 (p=0.89, WT: n=7, *rasa1-/-*: n=9, N=3). **E:** Diagram illustrates where subintestinal venous plexus (SIVP) images were taken and how plexus outgrowth was measured, perpendicularly from the DA to the ventral-most aspect of the SIVP. **F-I:** Confocal microscopy of wildtype (F, G) and *rasa1-/-* (H, I) on *Tg(flk:EGFP)* shown in black at 58hpf and 76hpf of the same embryos while the subintestinal venous plexus (SIVP) expands over the yolk sac. **J:** No change was seen in SIVP outgrowth at either timepoints (58hpf, WT: 190.2µm±36.0 and *rasa1* mutant at 179.1µm±41.8, p=0.45. 76hpf, WT: 212.6µm±36.6 and *rasa1* mutant SIVP at 232.0µm±43.6p=0.20, WT: n=17, *rasa1-/-* : n=12, N=3, paired t-test). Error bars represent ±SD.

**Figure 5.**
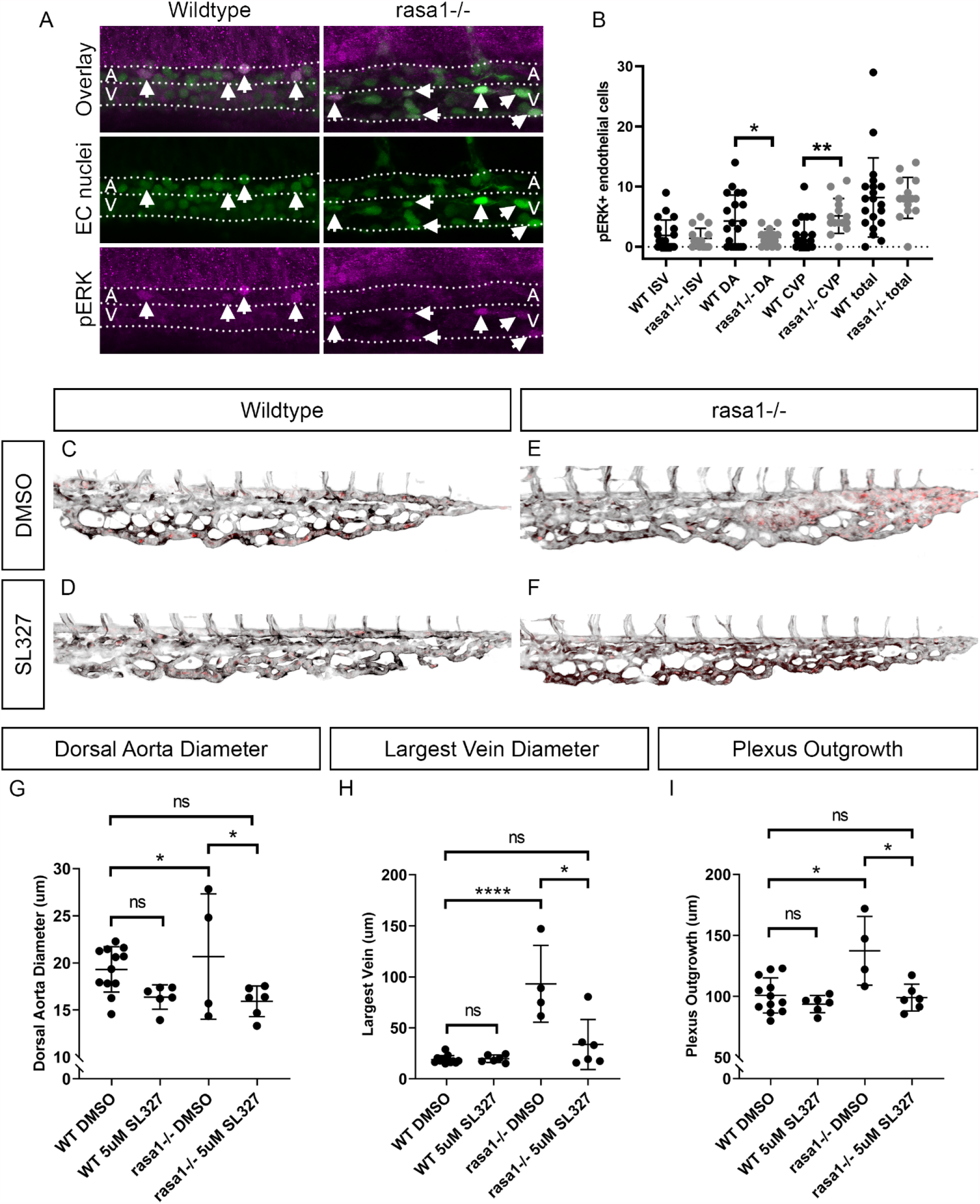
Ectopic venous activation of pERK in *rasa1* mutants may drive AVM formation, with cavernous malformations in *rasa1* mutants rescued by MEK/ERK inhibition. **A-B:** pERK antibody staining revealed and increase in pERK in the vein (WT: 2.0cells±2.6, *rasa1*-/-: 5.1cells±2.9, p=0.0020) and a decrease in the DA (WT: 4.3cells±4.3, *rasa1*-/-:1.6cells±1.4, p=0.03) with no change in ISVs (WT: 1.9cells±2.6, *rasa1-/-*: 1.4cells±1.7, p=0.52) (WT: n=20, *rasa1-/-:* n=15, N=3, unpaired t tests). **C-F:** Confocal images of wildtype and *rasa1* mutants on *Tg(flk:EGFP;gata1a:dsRed)* when treated with DMSO or SL327. Black is the *flk:EGFP* endothelium, red is *gata1a:dsRed* red blood cells. **G:** Quantification of dorsal aorta diameter in wildtype (WT) and *rasa1* mutants treated with DMSO and SL327 (WTDMSO vs. *rasa1-/-*DMSO: p=0.90, WTDMSO vs. *rasa1-/-*SL327: p>0.13, N=1, one-way ANOVA, Sidak’s multiple comparisons). **H:** Rescue of largest vein diameter is seen in *rasa1* mutants treated with SL327 (WTDMSO vs. *rasa1-/-* DMSO: p<0.0001, WTDMSO vs. *rasa1-/-*SL327: p=0.36, N=1). **I:** Plexus outgrowth is rescued by SL327 treatment of *rasa1* mutants treated with SL327 (WTDMSO vs. *rasa1-/-* DMSO: p=0.0013, WTDMSO vs. *rasa1-/-*SL327: p>0.99, N=1). P-values were calculated using a one-way ANOVA with Sidak’s correction for multiple comparison for SL327 experiments. Error bars represent ±SD.

Since MEK/ERK signaling is implicated in artery specification, we used in situ hybridization to assess if initial artery and vein specification were normal (Figure 3-figure supplement 1). Both arterial *ephrinb2a* and venous *ephb4a* expression at 30hpf appear identical in wildtype and *rasa1* mutants, suggesting early arteriovenous specification is unaffected (Figure 3-figure supplement 1).

To test whether venous pERK activation is important in AVM formation and if we could block AVM development, we applied 5µM SL327 (a MEK1/MEK2 inhibitor) from 24-48hpf. The largest veins of wildtype embryos treated only with DMSO vehicle control were 18.9µm±3.9 in comparison to *rasa1* mutants that measured 93.2µm±37.7 (p<0.0001, Figure 5C, E, H). MEK inhibition resulted in rescue of the enlarged vessels in *rasa1-/-* to a caliber indistinguishable from wildtype with DMSO (*rasa1*-/-_SL327_: 33.7µm±24.4, p=0.36, Figure 5C, F, G) and significantly smaller than DMSO-treated *rasa1* mutants (93.2µm±37.7, p<0.0001, Figure 5E-F, H). There is no significant change in DA diameter with DMSO treatment (p=0.90) or SL327 treatment in wildtypes or *rasa1* mutants (p>0.99, Figure 5C-G). We also tested whether the maximal migration distance is changed. We find that there is no significant difference in maximal distance of the plexus between vehicle treated wildtypes (100.8µm±14.4) and MEK inhibitor treated *rasa1* mutants (99.0µm±11.0, p>0.99, Figure 5C-F, I). Taken together we show that while artery-vein identity is initially not changed, activation of venous pERK is enhanced in *rasa1* mutants. pERK activation in the vein is functionally important since inhibition of MEK/ERK activity prevents AVM formation.

## Discussion

### Zebrafish *rasa1* mutants develop arteriovenous vascular malformations downstream of artery-vein specification

In vivo models of AVM formation with a stereotypical location are rare. We report that *rasa1* mutant (*rasa1a*^*-/-*^ and *rasa1b*^*-/-*^) zebrafish develop AVMs in the region of the caudal venous plexus (CVP) where the dorsal aorta (DA) turns into a venous plexus bed before returning to the heart. The vascular malformation we observe is an AVM because it develops and subsumes both vessels. In humans, vascular lesions arise from tissues where RASA1 incurs a somatic second hit (Cai et al., 2018; Lapinski et al., 2018; Macmurdo et al., 2016), while our model is a full genetic knockout. Phenotypes of single zebrafish *rasa1a* and *rasa1b* mutants are mild and a double knockout is necessary to produce highly penetrant phenotypes. *rasa1* mutant zebrafish do not survive to adulthood, and similarly, no humans have been identified with homozygous loss of function (pLI=1.0 for RASA1 in GnomAD v2.1.1). Rather, somatic mutations are found in the localized lesions of *RASA1* heterozygous CM-AVM patients, making the lesions homozygous for RASA1 mutations.

In the zebrafish tail, the DA and CVP are molecularly distinct, but directly connected. We observe vascular phenotypes initiate in the vein of *rasa1* mutants. A key metric to distinguish effects on the artery and vein is diameter. *rasa1* mutant AVMs have a massively increased diameter over normal CVP vessels but no change is seen in the artery. Genetic establishment of the artery or vein program occurs early in development, and key markers such as EphrinB2a and EphB4 are differentially expressed as early as 20-24hpf in development (Damm and Clements, 2017; Ren et al., 2013; Swift et al., 2014; Thisse et al., 2001). However, we show that arteries and veins appear to be correctly specified at 30hpf using the markers *ephrinb2a* and *ephb4a*. Thus, the zebrafish *rasa1* AVM likely develops after specification of artery and vein. This is not surprising given that RASA1 and EPHB4 are known to physically interact (Kawasaki et al., 2014), and that Rasa1 would act downstream of EphB4 receptor expression, but interfering with signaling and venous differentiation. EphB4 is a critical player in venous identity. Human CM-AVM2 results from mutations in EPHB4, further lending evidence to these two proteins acting in the same pathway (Amyere et al., 2017). EphB4 plays an important role in the separation of vein from artery through interaction with the arterial ligand, EphrinB2 (Hamada et al., 2003; Wang et al., 1998). Even if artery and vein are correctly specified, loss of EphB4 downstream signaling through loss of Rasa1 may affect the maintenance of the venous fate, resulting in incomplete separation of artery and vein. Other genetic forms of AVM are with altered signaling downstream in the Ras pathway also change arteriovenous signaling. KRAS gain-of-function changes expression of Notch pathway genes involved in arteriovenous specification (*DLL4, NOTCH1, HES1* and *HEY2*) but without the disruption of upstream *EPHRINB2* and *EPHB4* expression (Nikolaev et al., 2018). We propose that Rasa1 is necessary for the maintenance of venous fate, and its loss leads to aberrant connections with the artery, abnormal angiogenesis of the vein and upregulated pERK signaling (Figure 6).

**Figure 6.**
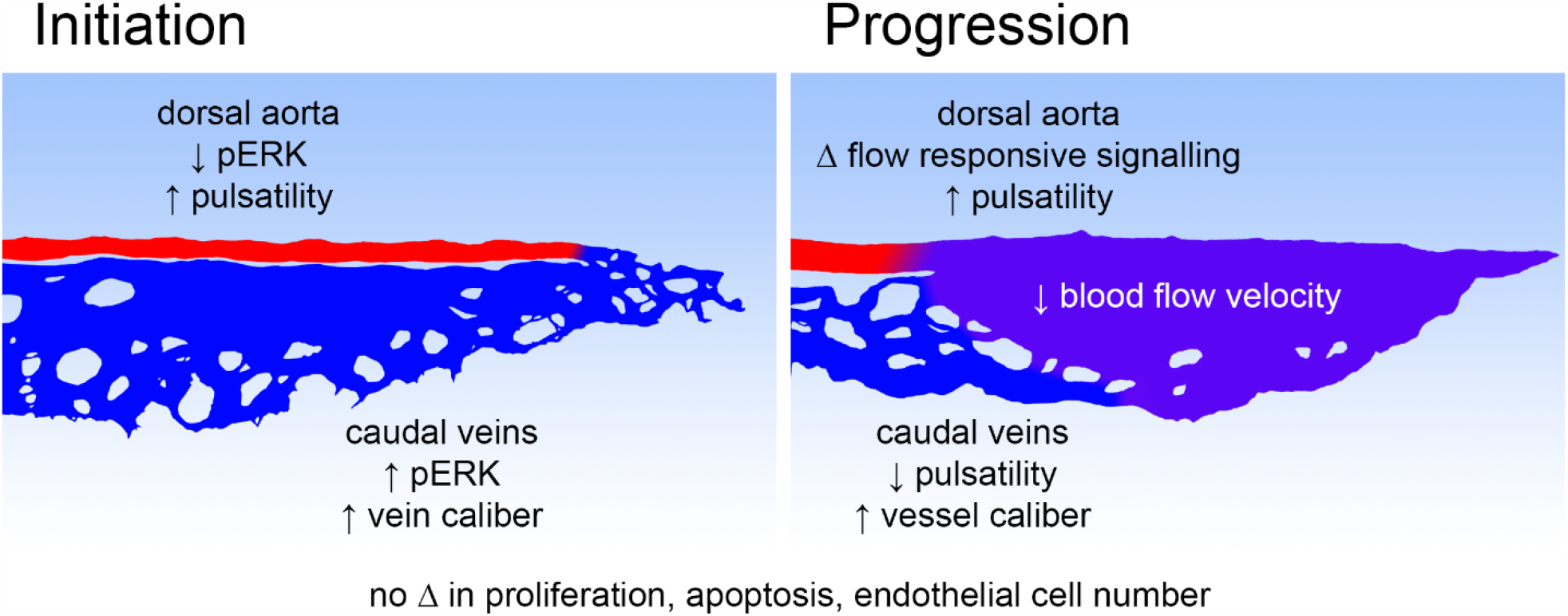
Model of *rasa1* AVM initiation and progression. During the initiation of *rasa1* AVM formation, there is an increase in venous pERK activation, vein caliber and pulsatility in the dorsal aorta. As the AVM progressively enlarges, there continues to be high pulsatility in the aorta, a drop in pulsatility in the caudal veins and well as slow blood flow velocity in the flow return. There are changes in flow responsive signalling including a decrease in *klf2a* expression. No changes in proliferation, apoptosis or endothelial cell number appear to drive the initiation or progression of *rasa1* AVMs.

### Ectopic pERK signaling in the vein of *rasa1* mutant AVMs

Loss of Rasa1 results in overactivation of Ras signaling and two potential downstream pathways, PI3K or MEK/ERK. Both pathways are drivers of AVMs in humans and animal models. The PI3K pathway is elevated in human and mouse models of HHT, and in human RASA1 vascular lesions (Alsina-Sanchis et al., 2018; Iriarte et al., 2019; Kawasaki et al., 2014; Ola et al., 2016). MEK/ERK signaling is upregulated in KRAS-caused brain AVM and in the Rasa1 mouse model as well as human RASA1 vascular lesions (Chen et al., 2019; Fish et al., 2020; Kawasaki et al., 2014; Lubeck et al., 2014; Nikolaev et al., 2018). Both MEK/ERK and PI3K/AKT/mTORC pathways are important in the specification of either artery (MEK/ERK) (Fischer et al., 2004; Hong et al., 2006; Lawson et al., 2001; Lawson et al., 2002; Shutter et al., 2000; Wythe et al., 2013) or vein (PI3K/AKT) (Chen et al., 2012; Fish and Wythe, 2015; You et al., 2005). Additional inhibitory loops between the two pathways to stabilize artery-vein identity (Hong et al., 2006). We observe a striking increased in venous MEK/ERK activation but not the adjacent dorsal aorta or intersegmental arteries of the trunk of *rasa1* mutants. While increases in pERK signaling have been previously seen in *Rasa1* mouse mutants, we are first to show the localization to the vein and that inhibition of MEK signaling prevents AVM formation. With the ectopic activation of MEK/ERK in the vein instead of the artery in *rasa1* mutants, venous programming may be disrupted, allowing for AVM formation. Over-activation of ERK in the vein may also upset angiogenesis and endothelial migration and permit the fusion of artery and vein (Shin et al., 2016; Srinivasan et al., 2009). Our data argue that the vein is particularly susceptible to perturbation of *rasa1* and that ectopic MEK/ERK signaling in the vein is implicated in the initiation of *rasa1* vascular malformations as inhibition of this signaling blocks lesion formation.

Our data suggest some mechanisms through which the cavernous AVM arises. Over the window of AVM formation, we did not observe changes in endothelial cell proliferation, cell death, or endothelial cell number. Though the role of endothelial cell proliferation varies between vascular malformations, our *rasa1* data are consistent with the lack of proliferation in *Kras* AVM models (Fish et al., 2020; Nikolaev et al., 2018). However our data contrasts with the Rasa1 mouse model, where disruption of angiogenesis results in hemorrhage and edema, apoptotic endothelial cells, vascular smooth muscle cells and cells in the lymphatic vessel valve leaflets are observed (Chen et al., 2019). In comparison, increased proliferation is seen in the vascular lesions of the CCM3 mouse model as well as *CCM3*-/- cell culture (Bravi et al., 2015; Malinverno et al., 2019) with mixed findings from HHT models (Corti et al., 2011; Rochon et al., 2016; Roman et al., 2002; Sugden et al., 2017; Tual-Chalot et al., 2014). Differences in proliferation across vascular malformations suggest proliferation is a feature of some but not all malformations, or potentially important only certain stages of malformation development (initiation or progression).

If cell number is not changed in AVM development, other mechanisms must be at play to drive pathological vessel enlargement. Aberrant rearrangement of endothelial cells within the CVP during a critical period of remodeling could result in malformations. One potential mechanism for lesion initiation could include a collapse of the stereotypical webbed plexus into a single large tube. Previous work in the *Rasa1* mouse model has implicated the failed export and deposition of collagen IV in the vascular basement membrane in the pathology of *Rasa1* mice, which would likely impact vessel stability and could lead to plexus collapse (Chen et al., 2019). Secondly, an enlarged vessel could be the result of disrupted intussusceptive angiogenesis, where vessels are split by the formation of intraluminal pillars. This form of remodeling is critical in the development of the CVP into its mature form (Karthik et al., 2018), though the molecular mechanism driving intussusceptive angiogenesis are poorly described. Finally, a localized increase in endothelial cell size that may be driven by the interplay of genetics and altered flow signaling could help drive the progressive enlargement of the vascular malformations. Our data are consistent with Rasa1 AVM formation involving changes at the level of cellular architecture, but further investigation is needed.

### Localization of *rasa1* AVMs in the venous plexus of zebrafish

A common mechanism of vascularization in vertebrates involves the formation of an immature web of equally sized vessels in a plexus, that is then remodeled over time to produce a mature branched vascular tree (reviewed in Heinke et al., 2012). The CVP follows this vessel formation mechanisms. The posterior caudal vein sprouts ventrally between 24 and 30hpf to form a temporary plexus (Choi et al., 2011). The plexus begins to remodel at approximately 48hpf, with most of the dorsal plexus vessels regressing into a single ventral vein. Why might this plexus be particularly prone to developing vascular malformations? Firstly, the CVP sits at the junction of the DA, a high-pressure vessel, which turns 180°, and splits into multiple smaller vessels providing a single transition point between high velocity, high pressure vessels and a lower velocity, lower pressure venous plexus. While laminar shar stress typically promotes endothelial health, the turbulent flow at the turn-around point may sensitize this location to deformation (Chappell et al., 1998; García-Cardeña et al., 2001; Mohan et al., 1997). Flow can be vasoprotective, preventing cerebral cavernous malformation (CCM) formation in the CCM1 zebrafish model and the CCM2 mouse model that normally develop lesions in lowly perfused venous capillaries in the brain (Li et al., 2019; Rödel et al., 2019). Flow vasoprotection may explain why vessel beds are differentially susceptible to developing different types of vascular malformations, perhaps including *rasa1* AVMs.

Secondly, the CVP is a hematopoietic niche during early development but as hematopoietic stem cells eventually hone to the kidney niche, it is no longer needed (Xue et al., 2017). Thus, this area may be less well stabilized than other areas of the vasculature as it is a temporary structure. It is possible that the active formation and remodeling of this vessel bed makes it particularly susceptible of malformation in early development. Third, since the malformations develop at the intersection between the DA and the CVP, this could predispose the region to form malformations with any perturbations of arteriovenous specification or maintenance.

### Vessel enlargement in *rasa1* mutants leads to flow abnormalities

The AVM in *rasa1* mutants impacts flow velocity and pulsatility without impacting heart rate. At 30hpf, the velocities in the caudal vein of *rasa1* mutants are substantially increased in comparison to wildtypes as the lesion is developing. By 48hpf, the high caudal vein velocity is dampened by the cavernous malformation. We were also interested to find that velocity extremes are impacted by presence of the AVM. By 48hpf, the DA in mutants experiences higher maximal velocities and lower minimum velocities than wildtypes and more pulsatile flow. These changes would greatly alter the forces detected by endothelial cells. Altered flow patterns change mechanosensing in the endothelium, and downstream molecular signaling through regulation of flow-responsive transcription factors such as Klf2 (Dekker et al., 2002; Lee et al., 2006; Parmar et al., 2006). Indeed, we find reduced *klf2a* expression in *rasa1* mutants. Klf2-driven flow responsiveness is critical for endothelial health, with mutation of Klf2 resulting in cardiac failure in mouse and fish models (Lee et al., 2006). Other vascular malformations show changes in Kfl2 expression. In CCM, the dysregulation of Klf2 suggests that changes in flow in vascular malformations can alter signaling and promote lesion progression (Li et al., 2019; Rasouli et al., 2018; Renz et al., 2015; Zhou et al., 2015; Zhou et al., 2016). Flow and flow-responsive signaling also play a role in zebrafish hereditary hemorrhagic telangiectasia (HHT) AVM models, with the development of AVMs being dependent on flow and impaired polarization seen in flow-deficient endothelial cells, resulting in larger cells and consequently larger caliber vessels (Corti et al., 2011; Sugden et al., 2017). *klf2a* expression alterations begin early in a *rasa1* mutant model, suggesting that altered flow may play a role in the progression of vascular lesions and the enlargement of malformations.

Our experiments are conducted at a stage prior to the recruitment of vascular mural cells to the aorta (or vein). Thus, the malformation develops in the absence of external stabilization. Since *rasa1* mutants have severe edema at later stages, it is not possible to test if recruitment of smooth muscle cells would help reduce pulsatility in the aorta at later timepoints (Ando et al., 2016; Stratman et al., 2017). The DA is more proximal to the heart and would be subject to more force from heart contractions. By 48hpf, the pulsatility across the DA and ventral vein becomes more consistent. In contrast, in *rasa1* mutants, the DA pulsatility remains elevated relative to the ventral vein over both timepoints and there is a significantly higher drop in pulsatility from the DA to the ventral vein. The cavernous AVM vessels acts as a damper on pulsatility, impacting flow velocity and pulsatility both up and downstream of the malformation, which would likely reduce signals for mural cell recruitment at a later stage. Changes in pulsatility from the cavernous malformation may also impact signaling that is specifically responsive to pulsatile flow (Lara et al., 2013; Shepherd et al., 2009), meaning that both changes in velocity and pulsatility could contribute to the progression and enlargement of a vascular lesion (Figure 6). Pharmacological reduction of blood pressure (and presumably shear stress) through propranolol appears to prevent the development of vascular lesions in human CCM (slow flow lesion) and promotes the resolution of infantile hemangioma (fast flow lesion) (Léauté-Labrèze et al., 2015; Li et al., 2021; Oldenburg et al., 2021; Reinhard et al., 2016). Lowering shear stress may have a protective effect against the de novo formation of vascular lesions as well as promoting the remodeling of a pre-existing lesions. Future investigations into the role of flow in the progression of fast and slow flow vascular malformations may offer further insights.

The genetic *rasa1* mutant zebrafish model has helped us focus on the AVM component of the CM-AVM disorder, highlighting how MEK/ERK signaling, arterio-venous signaling, blood flow and pulsatility interact to correctly separate artery and vein during development. We have shown the development of cavernous vascular malformations in the tail plexus and specifically implicate the ectopic activation of venous MEK/ERK signaling in their initiation. Perturbed blood flow and pulsatility from vessel malformations likely also contribute to the progression of the lesion as downstream flow responsive signaling is altered.

## Materials and methods

### Zebrafish husbandry and fish strains

All experimental procedures were approved by the University of Calgary’s Animal Care Committee (Protocol AC17-0189). Zebrafish embryos were maintained at 28.5°C and in E3 medium (Westerfield, 1995).

Transgenic lines used include: *Tg*(*kdrl:mCherry*)^*ci5*^ (Proulx et al., 2010), *Tg*(*flk:GFP*)^*la116*^ (Choi et al., 2007), *Tg(gata1a:dsRed)*^*sd2*^ (Traver et al., 2003).

*rasa1a*^*ca35*^ and *rasa1b*^*ca59*^ mutants were generated using CRISPR-Cas9 mutagenesis, following the methods outlined in Gagnon et al. 2014. Briefly, a 20-mer target with T7 promoter and constant Cas rev oligo were ordered (IDT) and annealed to synthesize sgRNA through in vitro transcription with MAXIscript T7 Transcription kit (Ambion, Cat. No. AM1312). *rasa1a*^*ca35*^ and *rasa1b*^*ca59*^ target sequences are listed in Supplemental Table 1. *rasa1a* ^*ca35*^ was created with the injection of a stop cassette, which was not injected in the creation of *rasa1b*^*ca59*^ allele.

Embryos were injected at the 1-cell stage with 1uL (∼200 ng/μl) sgRNA and 1uL 300 ng/μl nls Cas9 mRNA (and 1uL 10 μM stop codon cassette oligonucleotide in the case of *rasa1a*^*ca35*^). P0 injected embryos were raised and outcrossed and F1 embryos were screened from mutations. Mutant alleles were cloned and sequenced from genomic DNA.

### Genotyping

Genomic DNA (gDNA) was extracted from whole embryos as described in “PCR Sample Preparation” from ZIRC protocols (https://zebrafish.org/wiki/protocols/genotyping). PCR was performed with primers listed in Supplemental Table 1 and visualized on an agarose gel.

### Drug Treatments

Embryos were dechorionated before drug treatment at 24hpf and treated until 48hpf. SL327 (Sigma S4069) stock solution at 5mM concentration was heated to 65°C for at least 20 min prior to dilution to 10µM in E3 (dosage similar to Shin et al., 2016 at 15 µM at 20-30hpf). SL327 and DMSO control treatments were performed in a 24-well plate, with approximately 20 embryos per well (Supplemental Table 2).

### In situ hybridization

In situ hybridization for *ephb4a, ephrinb2a* and *klf2a* were performed as previously published (Lauter et al., 2011) with some modifications. Pre-hybridization and probe hybridization were performed in 50% formamide hybridization buffer (50% formamide, 5xSSC, 5 mg/mL torula yeast RNA, 50 µg/mL heparin, 0.1% Tween-20 in water) with 5% dextran sulfate. Embryos were washed 2x 5 mins with 50% formamide, 2xSSC, 0.1% Tween-20 at 60°C, 15 mins with 2xSSC at 60°C, 2x 30 mins with 0.2xSSC at 60°C and blocked with 10% non-specific sheep serum (NSS) in PBT for 1h. All ISH were performed with anti-digoxigenin FAB fragments conjugated with alkaline phosphatase in 10% NSS/PBT and probe detection was performed with NBT/BCIP were diluted in NTT (100 mM Tris (pH 9.5), 100 mM NaCl, 0.1% Tween 20 in water). Once the reaction was finished, embryos were fixed for 15 mins in 4% PFA and cleared in glycerol overnight before imaging.

### Antibody Staining

Phospho-p44/42 MAPK (ERK1/2) (Thr2020/Tyr204) antibody (Cell Signaling, #9101) staining was performed as previously published (Randlett et al., 2015).

### Confocal Microscopy

Zebrafish were mounted on glass bottom petri dishes (MatTek, Ashland, MA, Cat. No. P50G-0-30-F), using 0.8% low melt agarose (Invitrogen (Carlsbad, CA) 16520-050) dissolved in E3 fish medium. Confocal imaging used for vessel measurements and hematocrit were obtained using a Zeiss LSM 700. All images were obtained with the 488 nm and 555 nm lasers, with a slice interval of 1-3 μm with a 20X (NA 0.8) objective. Embryos imaged to characterize vessel morphology were anesthetized with 0.004% tricaine methanesulfonate (Sigma, A5040), whereas embryos imaged for hematocrit calculations were not anesthetized. Timelapse stacks were collected on a Zeiss LSM 700 at an interval of 15 minutes for 6 hours from 24hpf through 30hpf.

### 3D modelling of confocal images of tail vessels

For modelling vessels with Simpleware ScanIP (Synopsys, 2018), high resolution confocal images were captured with using AiryScan Fast imaging on a Zeiss AiryScan LSM880 confocal microscope with an Apo 40xW (NA 1.1) objective using the Argon multiline laser for 488 nm excitation and the DPSS 561nm laser for 555 nm excitation. Slice intervals for these images were 0.25µm. A mask was created from the image and further refined utilizing the Gaussian filter for smoothing, island removal for RBC artefact removal, and flood fill for ensuring the model was contiguous.

### Image analysis for vessel morphology

Images were processed using ImageJ/Fiji. Vessel diameters were measured at positions where there were no other vessels directly connecting to the vessel being measured. The vessel diameter was measured from the external diameter of the endothelium for the dorsal aorta, and internal diameter for the caudal veins due to the complex nature of this vessel bed. Three measurements were obtained across the dorsal aorta and averaged to obtain an average dorsal aorta diameter. Vein enlargement was measured as the internal diameter of the largest vessel in the caudal venous plexus. Vessel enlargement was designated as a vessel >1.5x the average wildtype vessel. CVP and SIVP outgrowth was measured perpendicularly from the ventral side of the DA to the ventral most aspect of the CVP/SIVP. Outgrowth speed was calculated from outgrowth measurements of the same embryos at two timepoint, divided by the time between imaging.

### Velocity, heartrate and pulsatility measurements with MicroZebraLab

Videos were taken of embryos mounted in low melt agarose without tricaine at 10x magnification and 120 fps using the MicroZebraLab apparatus created by Viewpoint Life Sciences Inc (ViewPoint Behaviour Technology, n.d.). Videos were analyzed using the Zebrablood program. Videos of the heart were used to measure heartrate over a minimum of 30 seconds. The average velocity across the vessel diameter, *ū*, was also measured. The pulsatility index (PI), which quantifies variation in blood velocity due to the heartbeat, was calculated with equation 6. The drop in PI between the dorsal aorta and ventral vein was calculated using equation 7.

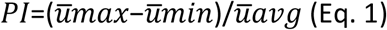

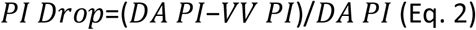

### Velocity heatmaps

Heatmaps of the velocities along the flow path in zebrafish embryos were created by using Microsoft Excel’s built-in conditional formatting and image processing via PaintTool SAI (SYSTEMAX Software Development), referencing an overlaid still image from MicroZebraLab detailing the boundaries of the measured region. The flow parameters were measured at 21 – 27 different locations per embryo, depending on flow path complexity and whether quality data could be obtained in the determined locations. Average velocity heatmaps were generated using the mean velocities from multiple embryos at predetermined positions across the dorsal aorta and caudal venous plexus.

### Statistics

GraphPad Prism8 was used to carry out all statistics. Unpaired two-tailed t-tests were used for two group comparisons and one-way ANOVAs for multiple comparisons with p-values from Sidak’s multiple comparisons reported unless otherwise indicated. Paired t-tests were used when analyzing data for vessel measurements over time, or velocity measurements from the same embryo. Velocity and pulsatility data were graphed as normalized to baseline measurements unless otherwise indicated. The Chi-squared test was used for *rasa1-/-* survival. Multiple t-tests with correction for multiple comparisons using the Holm-Sidak method were used for positional velocity and pulsatility data with adjusted p-values being reported. All data are represented as mean ± standard deviation (SD). All statistical analysis used p-values of 0.05 as a cut-off for significance (p<0.05=*, p<0.005=**, p<0.0005=***).

## Acknowledgements

This study was funded by University of Calgary Cumming School of Medicine, Canadian Institutes of Health Research and Faculty of Graduate Studies studentships to JGW, a Grant in Aid from the Heart and Stroke Foundation of Canada (G-16-00012741) and CIHR Project grant funding (PJT-168938) to SJC and NSERC Discovery funding to KR. We would like to thank the Childs lab members, including Dr. Jae-Ryeon Ryu, Dr. Thomas Whitesell, Dr. Charlene Watterston, Nabila Bahrami (MSc) and Dr. Suchit Ahuja who gave thoughtful feedback throughout this project and in reviewing the manuscript.

## Competing interests

No competing interests to declare.

**Supplemental Table 1.**
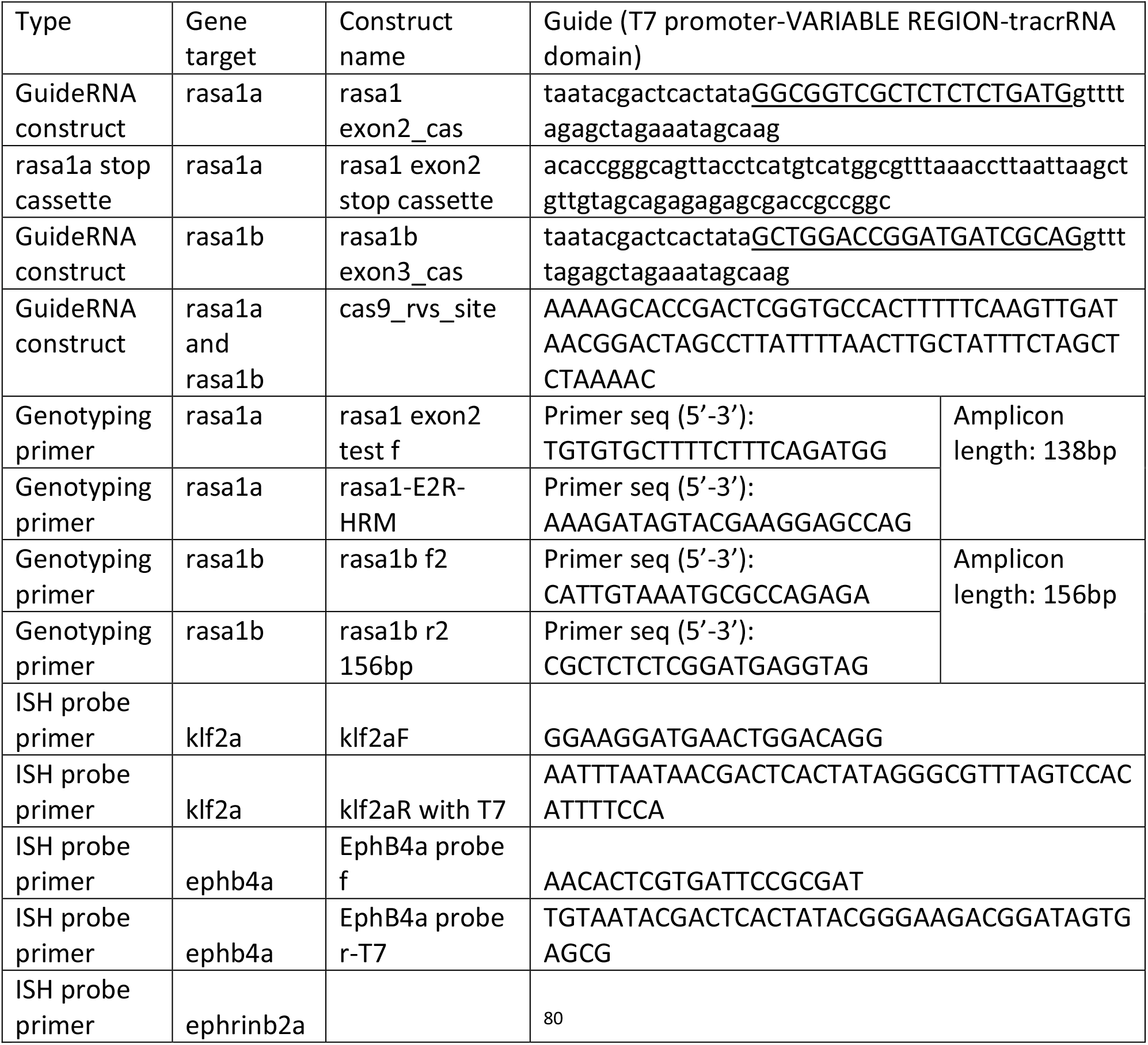
Constructs for GuideRNA, genotyping and ISH.

**Supplemental Table 2.**
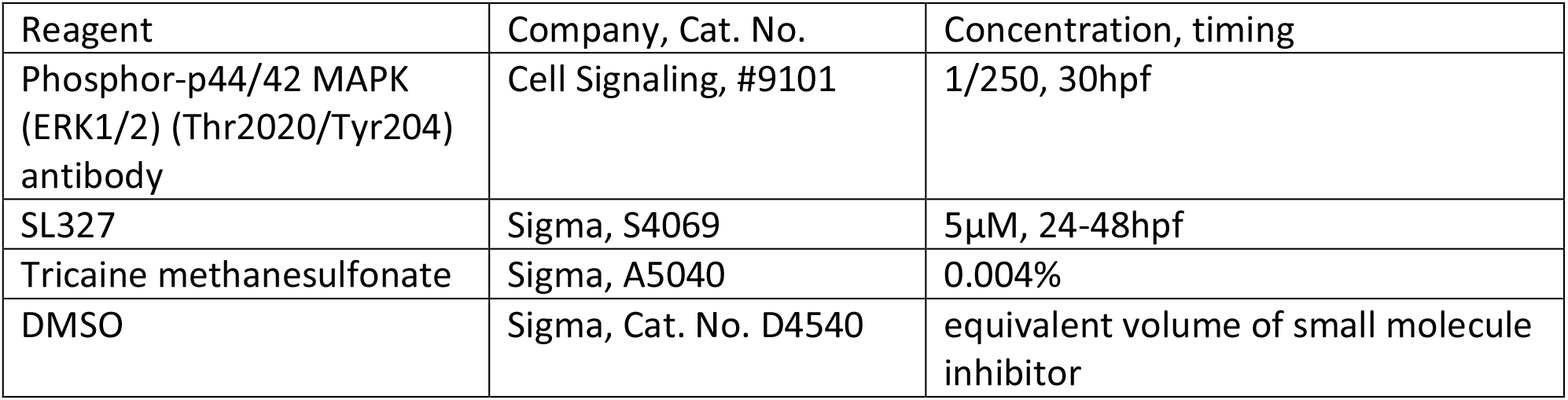
Source and concentration of reagents

| Reagent | Company, Cat. No. | Concentration, timing |
| --- | --- | --- |
| Phosphor-p44/42 MAPK (ERK1/2) (Thr2020/Tyr204) antibody | Cell Signaling, #9101 | 1/250, 30hpf |
| SL327 | Sigma, S4069 | 5μM, 24-48hpf |
| Tricaine methanesulfonate | Sigma, A5040 | 0.004% |
| DMSO | Sigma, Cat. No. D4540 | equivalent volume of small molecule inhibitor |

---

Video 1. High speed video imaging of blood flow through the dorsal aorta and caudal venous plexus of a laterally mounted wildtype embryo at 30hpf.

Video 2. High speed video imaging of blood flow through the dorsal aorta and caudal venous plexus of a laterally mounted *rasa1-/-* embryo at 30hpf.

Video 3. High speed video imaging of blood flow through the dorsal aorta and caudal venous plexus of a laterally mounted wildtype embryo at 48hpf.

Video 4. High speed video imaging of blood flow through the dorsal aorta and caudal venous plexus of a laterally mounted *rasa1-/-* embryo at 48hpf.

Video 5. Timelapse confocal imaging of the developing caudal venous plexus of a laterally mounted wildtype *Tg(kdrl:mCherry)* embryo from 24-30hpf. Imaging was performed with 15 min intervals and shown at 2fps.

Video 6. Timelapse confocal imaging of the developing caudal venous plexus of a laterally mounted *rasa1-/- Tg(kdrl:mCherry)* embryo from 24-30hpf. Imaging was performed with 15 min intervals and shown at 2fps. Note the cavernous AVM at the posterior of the tail, partially filled with stagnant blood.

## Notes

### Competing Interest Statement

The authors have declared no competing interest.

